# CH•••S hydrogen bonds drive molecular recognition of ergothioneine by the microbial transporter

**DOI:** 10.1101/2025.07.28.667264

**Authors:** Katherine A. Legg, Giovanni Gonzalez-Gutierrez, Katherine A. Edmonds, Philip G. Shushkov, David P. Giedroc

**Affiliations:** Department of Chemistry, Indiana University; Bloomington, IN, USA; Department of Molecular and Cellular Biochemistry, Indiana University, Bloomington, IN, USA

## Abstract

Many bacteria harbor an ATP-binding cassette (ABC) transporter named EgtU specific for the human dietary antioxidant and 2-thioimidazole-containing low molecular weight thiol ergothioneine (ET). How the solute binding domain, EgtUC, discriminates among ET and other similar molecules is unknown. Here, we use a “chimeric” mutagenesis strategy and two distantly related EgtUCs from *Streptococcus pneumoniae and Helicobacter pylori* to show that a suite of EgtUC alkyl CH•••S hydrogen bonds to the ET thione S atom are central determinants of molecular recognition. Small perturbations in CH•••S distance and angle give rise to sharply attenuated transport-competent ET-bound “closed” state lifetimes and increased motional disorder in the binding pocket, not around the S atom itself, but distally in weakening NH•••O hydrogen bonds. This is, to our knowledge, the first work to describe the impact of alkyl CH•••S H-bonding in a biological protein-ligand complex in water.

## Introduction

Cell-abundant low molecular weight (LMW) thiols maintain the reducing environment of the cytoplasm of bacterial cells and the cytosol of eukaryotic cells, and include the ubiquitous tripeptide glutathione (*1*). Bacteria unable to synthesize or import glutathione synthesize other thiols, including bacillithiol and mycothiol found in some Firmicutes and Actinomycetes, respectively (*2, 3*). These abundant LMW thiols scavenge endogenous or exogenous reactive oxygen and nitrogen species (ROS, RNS), creating thiol disulfides that are subsequently enzymatically reduced, thus maintaining redox balance (*4*). LMW thiols in bacterial pathogens play critical roles in oxidative, reductive, electrophile and transition metal stress responses in the infected host (*5-8*).

Many bacteria utilize more than one LMW thiol capable of playing distinct functional roles (*5-10*). Previous work reveals that the LMW thiol *L*-ergothioneine (ET), a potent dietary antioxidant and cytoprotectant of therapeutic value in humans (*11-14*) is a cell-abundant LMW thiol that is imported by many bacterial phyla via an ATP-binding cassette (ABC) transporter (*15*) which we denote EgtU (*16, 17*). *L*-ergothioneine (EGT or ET) is trimethylamine (betaine) derivative of histidine with a sulfur atom installed at the imidazole C2 position, with the thione tautomer dominant at neutral pH (*18*). In vertebrates ET becomes broadly bioavailable in tissues and cells via the action of the ubiquitously expressed ET transporter ETT (*19-21*). The primary role of ET in vertebrates appears as a nutritional antioxidant, in particular as a one-electron scavenger of myriad ROS, including H_2_O_2_, superoxide anion and ^1^O_2_ (*22-24*). Oxidized forms of ET (ESSE, 5-oxo-ET, ET sulfonate) can be recycled by glutathione reductase in the presence of glutathione (*23*). Other functions in vertebrates are possible; a recent report suggests that ET provides an important source of bioactive sulfur, which upon oxidation leads to the formation of reactive sulfur species (RSS) that enhance energy generation and lifespan via enzyme persulfidation (*25*).

The physiological functions of ET in bacteria are far less well-understood. ET is biosynthesized by select fungi and bacteria from histidine, notably *Mycobacterium tuberculosis*, where it may function as a scavenger of ROS and is a known virulence determinant in infections (*26, 27*). The gut commensal *Lactobacillus reuteri* imports ET through an as yet-uncharacterized EgtU-like transporter and may directly impact gastrointestinal levels of ET (*28*); other microbes are known to utilize a histidinase-like enzyme to metabolize ergothioneine to thiourocanate, which is subsequently degraded to produce hydrogen sulfide (*29, 30*). In *Helicobacter pylori*, EgtU provides a competitive colonization advantage in infected mice; further, this work reveals that ET is metabolized by bacteria for protection and thus may play a yet unappreciated role in gut bacteria, both commensals and pathogens alike (*17, 31*).

The emerging importance of ET in bacterial physiology motivates a detailed understanding of molecular recognition of this key metabolite by the microbial transporter. EgtU is a heterotetramer of two subunits, EgtUA and EgtUBC. EgtUA is the ATPase that reversibly associates with EgtUBC, a fusion protein of the N-terminal EgtUB transmembrane domain, which forms the channel for ET translocation, and the C-terminal EgtUC domain, which is necessary and sufficient for ET molecular recognition (*15*). In *Streptococcus pneumoniae* (*Sp*) EgtUC binds ET as a 1:1 complex with a *K*_a_ of ≈2.0×10^7^ M^-1^. The crystallographic structure of *Sp*EgtUC in the ligand-bound form reveals two subdomains, D1 and D2, connected by two 10-residue strands, with the ligand bound in the cleft between subdomains (Fig. 1A) (*32*). The quaternary amine is oriented toward the hinge as it is in other betaine compound transporters; the thione sulfur atom, as part of a C-S double bond is close to the opening of the binding pocket (Fig. 1A). Detailed biophysical studies (*33*) and two structures of the ligand-free solute binding domain of EgtU from *H. pylori* (denoted *Hp*EgtUC in this work), and *Listeria monocytogenes* (formerly known as BilEB) (*17, 34*), suggest an open-to-closed, rigid-body 54° rotation of one subdomain relative to the other upon ET binding to *Sp*EgtUC. Strikingly, unlike vertebrate ETT, EgtUC is exquisitely selective for *L-*ET relative even to *L*-hercycine (HER), binding ET some 10^4^-fold more tightly. HER lacks only the thione sulfur of *L*-ET and can be a marker for oxidation of ET in cells (*35*). Glycine-betaine and *L*-histidine also bind very weakly (*16*).

**Figure 1.**
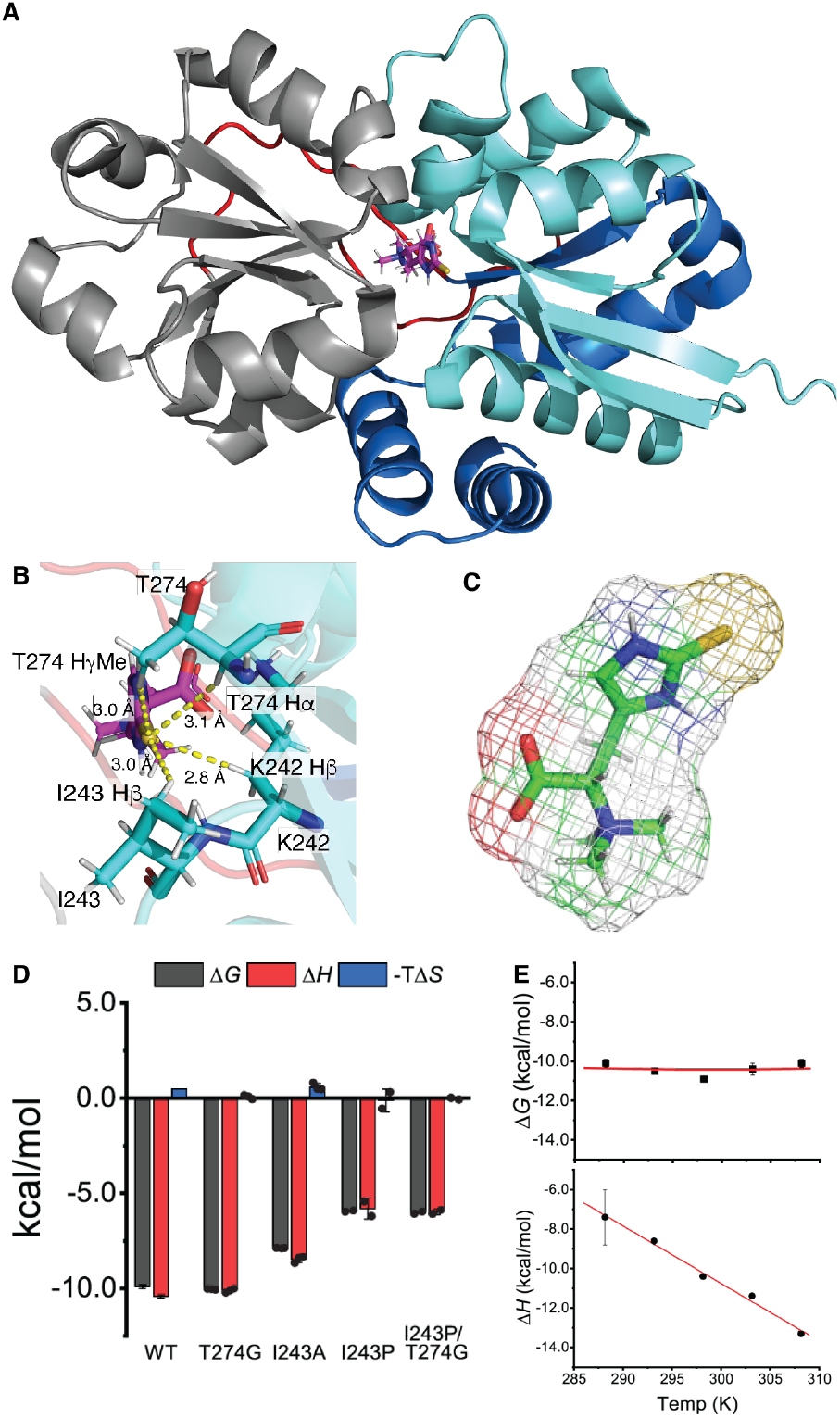
*Sp*EgtUC binding pocket, focusing on C-H•••S H-bond donating residues targeted for mutagenesis studies. (**A**) Crystal structure of *Sp*EgtUC bound to ET (PDB: 7TXK) (**B**) C-H•••S hydrogen bond donating residues of interest in this study. (**C**) Spacefill structure representation of *L*-ET. (**D**) Energetics of ET binding by *Sp*EgtUC and mutants. WT *Sp*EgtUC plotted as previously described (*16*). (**E**)Temperature dependence of free energy (top) and enthalpy (bottom) of ET binding to WT *Sp*EgtUC, with the continuous lines a fit to a nonlinear model (*47*) (*top*; with Δ*C*_p_ fixed to the experimental value (bottom panel) and *T*_H_= 263.1 K and *T*_S_=298.9 K) and linear model (*bottom*), which assumes a *T*-independent Δ*C*_p_=*d*Δ*H*/*dT*. See fig. S3 and table S1 for representative titrations and compiled thermodynamic parameters.

A functionally critical and unique feature of the bacterial ET transporter is a suite of noncovalent interactions between *Sp*EgtUC and the thione sulfur atom of ET that we hypothesize meet the definition of alkyl CH•••S hydrogen bonds (Fig. 1B) (*16, 36*). In the work described here, we critically evaluate the energetic, dynamical and structural consequences of perturbing individual candidate CH•••S H-bonding interactions in *Sp*EgtUC in an effort to understand thione sulfur recognition at the molecular level. A comparison of our ET-bound EgtUC structure to the distantly related EgtUC from *H. pylori* (*17*) reveals that the two residues that make closest approach to the S atom are not conserved (I243 in *Sp*, P296 *Hp*; T274 *Sp*, G327 *Hp*). Here, we use a “chimeric” mutagenesis approach to show that the ET-binding site is differentially sensitive to amino acid substitutions. While the T274G mutant is effectively silent, the I243P substitution dramatically reduces the lifetime of the ET-bound “closed” state (by ≈10^3^-fold), not by perturbation of the CH•••S bonds, but via a destabilization of NH•••O H-bonds donated by K242 and R379 that electrostatically “clamp” the ET carboxylate group distal to the S atom. We conclude that these CH•••S H-bonds anchor the thioimidazole ring of ET in the binding pocket, thus providing a means to ensure specificity of EgtU for thione S-containing aromatic metabolites. This work represents to our knowledge the first significant attempt to explore the impact of CH•••S H-bonding in molecular recognition in a biological protein-ligand complex in water.

## Results

### CH•••S H-bonds impact ET binding affinity and thermodynamics

CH•••S H-bonds are often overlooked noncovalent interactions that, like conventional H-bonds involving a pair of more strongly electronegative atoms, are both polarizable and directional (*36, 37*). CH H-bond donors in synthetic host-guest systems in organic solvent tend to have a particular affinity for larger anions, including halides (X^−^) and HS^−^, and can originate with both aryl and alkyl CH donors (*38, 39*). The optimal interatomic alkyl CH•••S distance is ≈3.0 Å, well within the sum of the van der Waals radii of both atoms, with a modestly more acute alkyl CH•••S bond angle (≈160º) found upon survey of the Cambridge Structural Database (*36*), relative to the shorter distance (≤2.0 Å) and linear angular dependence of conventional NH•••(N/O) H-bonds (*39*). A more strongly polarized CH bond is expected to function as a stronger donor, with the trend in anticipated H-bond strength in proteins being aryl>Hα>methine>methylene>methyl (*36*). Given that the thione conformer of ET exhibits significant partial negative charge on the S atom (–0.49) (Fig. 1C), we reasoned that CH H-bond donors in EgtUC might play a critical role in molecular recognition of ET.

While a number of mutations in the EgtUC binding pocket, *e*.*g*., those in the aromatic cage (*40*) that interact with the betaine moiety, predictably and negatively impact the affinity of EgtUC for ET, the importance of candidate CH•••S hydrogen bonding interactions of the ET thione sulfur have not been previously investigated (*16*). We therefore chose to elucidate the thermodynamics of ET binding by *Sp*EgtUC mutants using isothermal titration calorimetry (ITC). Previous data reveal that the binding of wild-type *Sp*EgtUC to ET is enthalpically driven at 25 ºC with a significant Δ*H* of ≈ –10 kcal mol^-1^ and a very small unfavorable -*TΔS* (*16*). We first substituted the invariant K242 with Ser or Ala, each of which would retain an Hβ2-like CH•••S H-bond donor atom (Ser, methylene group; Ala, methyl group); however, both substitutions will also disrupt the electrostatic interaction with the ET carboxylate (*16*). As expected, both K242S and K242A EgtUC substitutions abrogate ET binding (Table 1), thus suggesting this electrostatic interaction overrides the presence of a candidate CH•••S H-bond here. In contrast, substitution of T274 with glycine (T274G), as found in *H. pylori* EgtUC (*17*), has no discernable impact on the ET binding affinity or underlying energetics in *Sp*EgtUC (Table 1, Fig. 1D, fig. S1). This reveals that the CHα•••S H-bond is energetically far more important that any CHγ•••S H-bond, a finding consistent with expectations from the polarizability of the C-H bond (*36*).

**Table 1.**
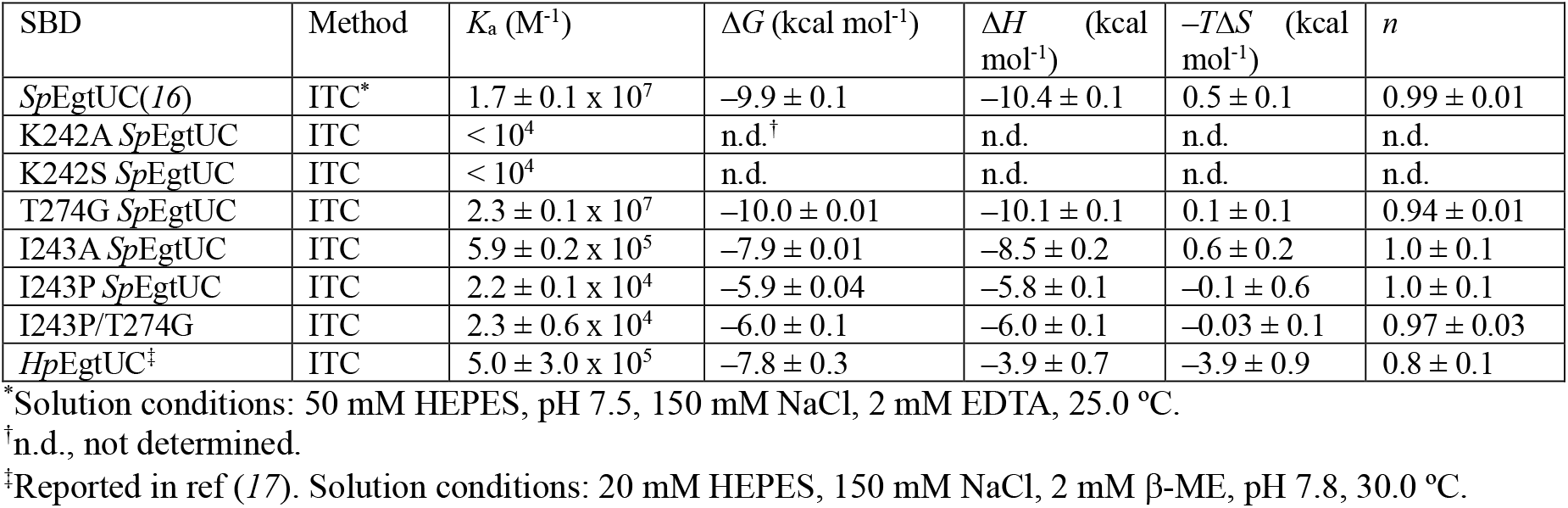
Thermodynamic parameters obtained for the binding of ET by *Sp*EgtUC and mutants.^a^.

Substitution of I243 to alanine reduces the affinity for ET by ≈30-fold, while substitution to proline, as in the *H. pylori* EgtUC sequence, reduces the affinity by ≈1000-fold (Table 1). The thermodynamic impact on ET binding by I243A EgtUC lies exclusively in the enthalpy term consistent with weaker noncovalent interactions in this mutant (Fig. 1D). Remarkably, I243P EgtUC, like wild-type EgtUC and other mutants, maintains a Δ*H* that is nearly identical to Δ*G*, with a negligible entropy (*T*Δ*S*) contribution (Table 1, Fig. 1D). This is further evidence that the impact of this highly destabilizing I243P substitution occurs as a result of local perturbations in noncovalent interactions rather than other contributors to the binding equilibrium, *e*.*g*., large changes in solvation. All mutants retain specificity for ET over HER with no binding of HER detected by ITC (fig. S2B-G) thus revealing specific elements of molecular recognition of ET are retained in this mutant. We also characterized the double mutant, I243P/T274G *Sp*EgtUC, in an effort to rescue the destabilizing effect of the I243P substitution in creating an *H. pylori*-like ET binding site on an otherwise *S. pneumoniae* EgtUC background. This double mutant is indistinguishable from the parent I243P *Sp*EgtUC (Table 1, Fig. 1D); this lack of rescue reveals that other as yet unknown *Hp*EgtUC-specific interactions are not recapitulated in this otherwise all *Sp* mutant. Further investigation of the thermodynamics of ET-binding by *Sp*EgtUC reveals a large negative change in heat capacity (Δ*C*_p_ of –0.29 kcal mol^-1^ K^-1^), with a nearly *T*-independent Δ*G* of binding (15–35 ºC) (Fig. 1E; fig. S3; Table 3, table S1). These findings are as anticipated from a significant decrease in solvent-accessible surface area (SASA) upon ET-mediated closure of the domain (Table 3) (*16, 41, 42*), which we estimate to be ≈14 residues in both the ET and I243P *Sp*EgtUCs (see below; fig. S4).

### Crystal structures of ET-bound mutant *Sp*EgtUCs are globally identical to the wild-type protein

We next sought to understand the impact of these substitutions on the suite of CH•••S and conventional NH•••N/O H-bonds in the EgtUC-ET complex. We obtained crystallographic structures for four ET-bound *Sp*EgtUC mutants (I243A, I243P, T274G, I243P/T274G) (Fig. 2, fig. S5) (see table S2 for crystallographic statistics) to a resolution of 2.1 Å or better with the respective structures (residues 233-506) solved by molecular replacement with the wild-type complex (*16*). The global structure of each mutant is essentially identical to WT *Sp*EgtUC, regardless of the change in binding affinity for ET. The trimethylamine cation binding “cage” is unchanged in all cases. In the T274G mutant, the CH•••S H-bond distances and angles are essentially wild-type (±0.1 Å in distance; ±10º in angle, Table 2, fig. S5B); this tracks with an ET affinity and underlying energetics that are indistinguishable from the wild-type domain. Thus, the γ-methyl group T274 makes no significant contribution to the ET-complex stability.

**Table 2.** C-H•••S H-bond lengths and bond angles in wild-type and mutant *Sp*EgtUCs.

| | | | 242 Hβ | | 243 H$\beta^{\P}$ | | 243 Hγ | | 243 H$\delta 1$ | | 274 H$\alpha^{\#}$ | | 274 Hγ Me | |
| --- | --- | --- | --- | --- | --- | --- | --- | --- | --- | --- | --- | --- | --- | --- |
|  | PDB Code | Resolution (Å) | Length (Å)<sup>†</sup> | Angle<sup>‡</sup> | Length (Å) | Angle | Length (Å) | Angle | Length (Å) | Angle | Length (Å) | Angle | Length (Å) | Angle |
| WT* | 7TXK | 1.8 | 2.8 | 177.2 | 3.0 | 125.8 | 2.9 | 130.7 | - | - | 3.1 | 152.1 | 3.0 | 152.4 |
| T274G | 9EGI | 2.0 | 2.7 | 176.8 | 2.9 | 123.9 | 2.9 | 121.6 | - | - | 2.9 | 156.8 | - | - |
| I243A | 9EGH | 2.1 | 2.5 | 153.1 | 3.2 | 133.1 | - | - | - | - | 3.1 | 167.2 | 3.6 | 145.8 |
| I243P | 9EGJ | 1.9 | 2.8 | 169.5 | 3.5 | 117.6 | 3.5 | 100.6 | 2.3 | 151.5 | 3.2 | 149.7 | 2.9 | 157.3 |
| I243P/T274G | 9O5F | 1.4 | 2.9 | 170.3 | 3.8 | 115.3 | 3.3 | 117.7 | 2.9 | 136.0 | 3.0 | 137.9 | - | - |
| *Hp EgtUC*<sup>§</sup> | 8DP7 | 3.4 | 3.4 | 163.2 | 6.0 | 63.6 | 3.6 | 113.6 | 2.9 | 150.9 | 3.0 | 136.8 | - | - |

\*PDB 7TXK, published previously (16).
<sup>†</sup>CH...S (hydrogen-sulfur) distance
<sup>‡</sup>C-H...S angle (180° to -180°)
<sup>§</sup>PDB 8DP7, published previously (17). Only distances $\leq 4.0$ Å are listed in this table, except where indicated.
<sup>\P</sup>This corresponds to Hβ
<sup>#</sup>This corresponds to H $\alpha 1$

**Figure 2.**
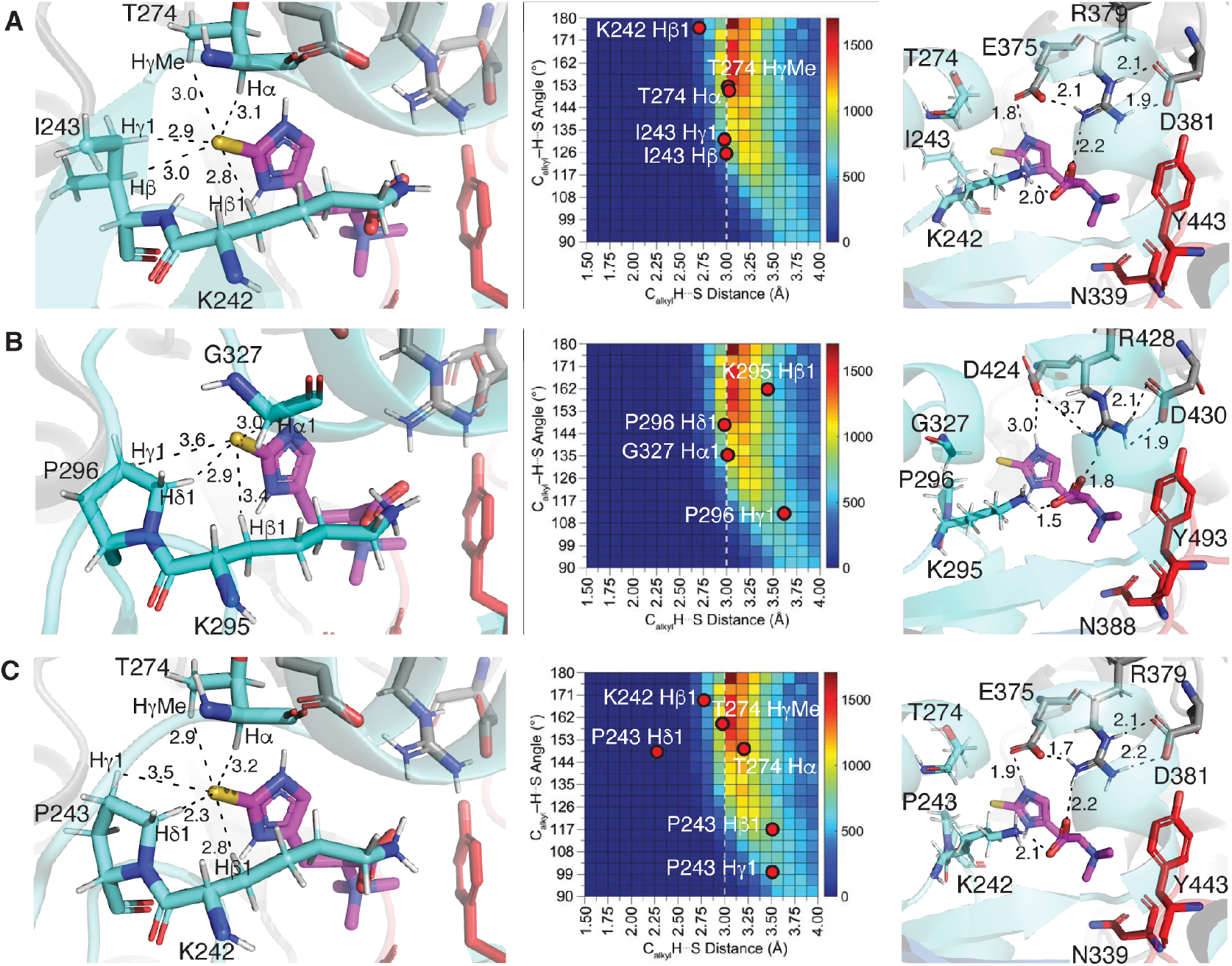
Crystallographic analysis of various EgtUC domains. *Left to right*, C-H•••S H-bonding interactions with H-S bond lengths noted, heatmap of a histogram of alkyl C-H•••S H-bond lengths and angles from an analysis of the Cambridge Structural Database (CSD) reproduced from reference (*36*) with experimental CH•••S H-bonds superimposed on this map, and N-H•••O hydrogen bonding interactions in (**A**) wild-type *Sp*EgtUC, (**B**) wild-type *Hp*EgtUC, and (**C**) I243P *Sp*EgtUC.

In the I243A structure, the most significant change is in the T274 γ-methyl protons, which have moved away from the sulfur atom by ≈0.5 Å (Table 2); however, this is not expected to impact the affinity, given the behavior of T274G *Sp*EgtUC. Thus, the poorer H-bond donating ability of the methyl vs. methine hydrogen in A243 vs. I243 must explain the reduced heat of binding on an otherwise wild-type structure (Table 1, Fig. 1D). In I243P EgtUC, P243 exists in the lower-energy *trans* configuration, thus ruling out the possibility that a *cis*-Pro in this position might broadly impact ET binding; the same is true of P296 in *Hp*EgtUC (*17*). In addition, interactions from the T274 side of the complex are identical. This suggests that small perturbations in CH•••S H-bonding distance and angle originating from residue 243 are responsible for the dramatically reduced affinity in this mutant. The Hβ methylene pair in *Hp*EgtUC is too far away from the sulfur atom, leaving the Hγ and Hδ hydrogens as the primary H-bond donors to the thione sulfur of ET (Table 2, Fig. 2B, left). While the Hβ of the I243P structure is oriented such that it is a similar distance from the thione sulfur as Hγ (Table 2, Fig. 2C, left), these interactions, appear less favorable in the I243P vs. WT *Sp*EgtUC structures (Fig. 2A, 2C, middle) (*36*). This collectively suggests that the degree of substitution of the CH donor at residue 243 has a major impact on ET binding affinity, subtly perturbing non-covalent interactions that strongly impact the Δ*H* of binding (Fig. 1D).

In addition to these small differences in alkyl CH•••S H-bonds among the *Sp*EgtUC mutants, it seemed possible that small changes in more conventional H-bond interactions might be present, thus impacting the energetics and affinity of ET binding. These H-bonding interactions (*16*) include an ET Nε2-H•••E375 Oε1 H-bond, and an electrostatic “clamp” around the carboxylate moiety of ET, involving K242 and R379. In addition, E375 and D381 stabilize this binding pocket. Strikingly, in each mutant, perturbations to these conventional hydrogen bonds are minimal overall (±0.2 Å in distance) in these crystallographic structures (Fig. 2, S3 right).

### NMR studies of I243A and I243P *Sp*EgtUCs reveal shorter closed-state lifetimes and weakened H-bonding

Given that ET binding is coupled to a large conformational change that buries ET at the D1-D2 interdomain interface (Fig. 1E; Table 3), we reasoned that perturbations in CH•••S H-bonds derived from residue 243 might have a significant impact on the lifetime (stability) of the ET-bound “closed” state. Previous NMR studies (*16*) reveal that *Sp*EgtUC exists in two conformations in solution: an open, ligand-free structure and the ET-bound closed state that are in slow chemical exchange on the NMR timescale (≈ms) (fig. S6). In addition, the apoprotein does not significantly populate the closed conformation in the absence of ET, consistent with ligand-mediated “induced fit” model (*16*). ^1^H,^15^N TROSY spectra of the EgtUC mutants I243A and I243P in the ligand-free state both closely resemble the wild-type spectra (fig. S7), with chemical shift perturbations localized to the site of the mutation, consistent with minimal changes in the overall structure of the domain.

**Table 3.** Change in molar heat capacity (Δ*C*_p_) and estimated change in solvent-accessible surface area (ΔSASA) upon ET binding to *Sp*EgtUC.

| $\Delta C_p$ (kcal mol<sup>-1</sup> K<sup>-1</sup>)* | ΔSASA<sub>ap</sub><br>(Å<sup>2</sup>)<sup>†</sup> | ΔSASA<sub>p</sub><br>(Å<sup>2</sup>)<sup>‡</sup> | ΔSASA<sub>MD</sub><br>(Å<sup>2</sup>)<sup>§</sup> | SASA<sub>ET</sub><br>(Å<sup>2</sup>)<sup>¶</sup> | #AA<sup>#</sup> |
| --- | --- | --- | --- | --- | --- |
| -0.292 | 648 | 1122 | 1160±66 | 255/451 | 19-20/14 |

\*Calculated from Fig. 1E.
<sup>†</sup>Estimated change in solvent accessible surface area (SASA), assuming all folded residues are apolar.
<sup>‡</sup>Estimated change in SASA, assuming all folded residues are polar.
<sup>§</sup>Computed change in SASA from molecular dynamics (MD) simulations
<sup>¶</sup>SASA of ET, calculated using PyMOL “get\_area” function. Ergothioneine was modeled using MolViewer (48). Second value is the GROMACS computed SASA value of ET with default parameters.
<sup>#</sup>Number of amino acids involved in the ET-induced folding of *SpEgtUC*. Average surface areas of polar and apolar ligands as reported (41). Second value is computed number of amino acids involved in the ET-induced folding of *SpEgtUC* from MD simulations (see fig. S4).

In striking contrast to WT *Sp*EgtUC, an ET titration of I243A EgtUC moves the ligand-bound complex into an intermediate chemical exchange regime, where most of the resonances that move are strongly line-broadened at sub-stoichiometric ET, revealing sharp resonances and WT-like chemical shifts only upon saturation of the complex (Fig. 3A, fig. S8A). These findings make the prediction that the I243P mutant complex will be in fast chemical exchange with the unbound conformation on the NMR timescale, and this is exactly what we find (Fig. 3B, fig. S8B). A global fit of these data to a 1:1 binding model reveals a *K*_a_ of ≈2.2 × 10^4^ M^-1^ (Fig. 3C), fully consistent with the calorimetric data (Table 1). If one further assumes that the association rate constant (on-rate) for ET binding is similar in all three cases and near diffusion-limited, *k*_on_≈10^8^ M^-1^ s^-1^ (both reasonable assumptions given the unobstructed ligand binding site in the apo state (*17*)), and a representative ^1^H chemical shift perturbation of ≈0.3 ppm (≈200 Hz at 600 MHz ^1^H frequency) (Fig. 3A-B), we estimate that the lifetime, τ (1/*k*_off_) is ≈5 ms for I243A *Sp*EgtUC (in the middle of the intermediate exchange regime for a representative nucleus). The τ for wild-type *Sp*EgtUC is then ≈150 ms, and for the weak-binding I243P mutant ≈150 µs. These thermodynamic and kinetic data strongly argue that perturbation of the ligand-binding pocket in the I243 substitution mutants significantly decreases the lifetime of the closed, transport-competent conformation.

**Figure 3.**
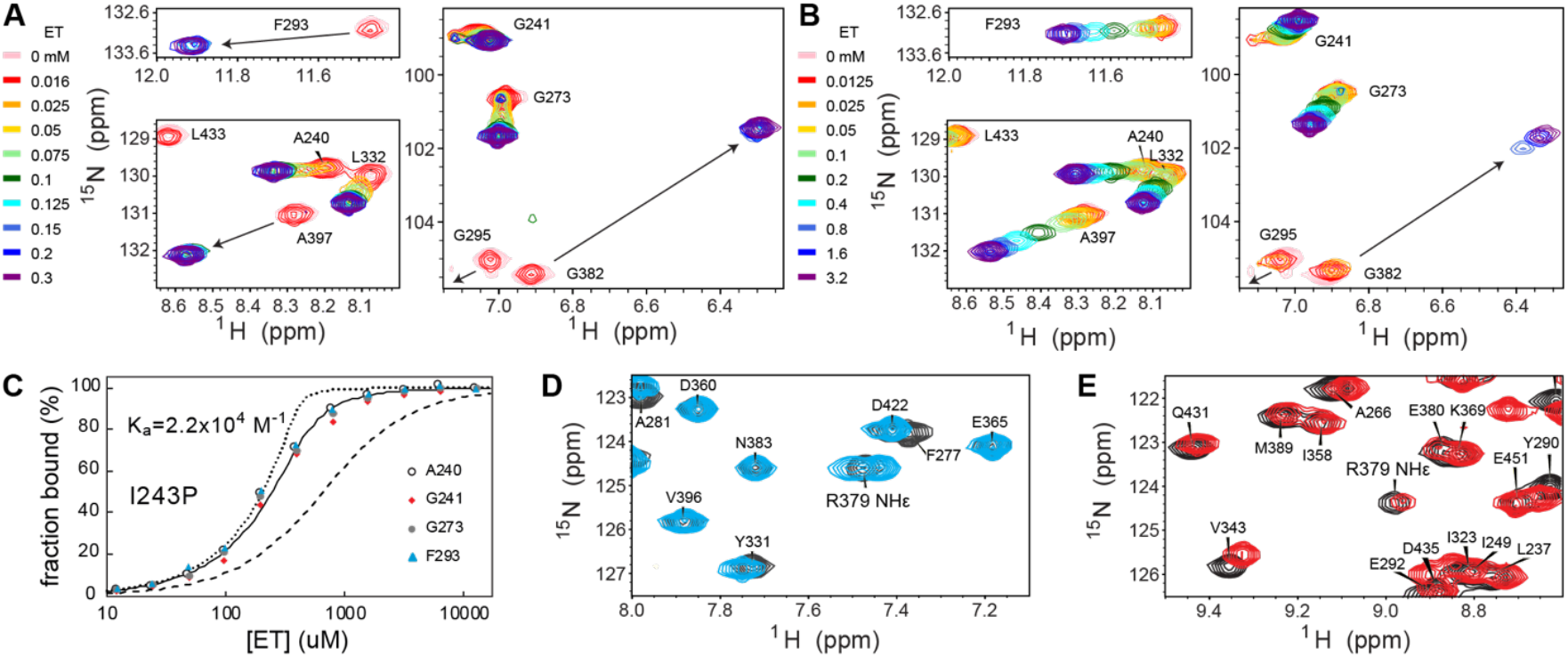
NMR of I243A and I243P *Sp*EgtUC. (**A**) Select regions of the ^1^H,^15^N TROSY spectra of ^15^N I243A SpEgtUC showing peak shifts of indicated residues upon titration with ET (see Fig, S6A for the full spectra). (**B**) Select regions of the ^1^H,^15^N TROSY spectra of I243P *Sp*EgtUC showing peak shifts of indicated residues upon titration with ET (see Fig, S6B for the full spectra). (**C**) Binding curve of ^15^N I243P *Sp*EgtUC titrated with ET calculated from chemical shift perturbations of the ^1^H,^15^N TROSY crosspeak of the indicated residues. The solid line shows a global fit of those four residues, with a *K*_a_ of 2.2 × 10^4^ M^-1^. Dashed and dotted lines show theoretical binding curves for binding 10-fold weaker and 10-fold tighter, respectively. (**D**) Select region of the spectrum of apo I243P (cyan) overlaid on the spectrum of apo WT (gray), highlighting the NHε cross peak of the R379 side chain. (**E**) Select region of the spectrum of ET-saturated I243P (red, 12.8 mM ET) overlaid on the spectrum of ET-bound WT (dark gray), highlighting the NHε peak of the R379 side chain.

Previous NMR experiments reveal that the chemical shift of the R379 NHε crosspeak moves substantially downfield (Δδ=1.5 ppm in ^1^H) and exhibits strongly attenuated motional disorder on the ps-ns timescale (^15^N-^1^H-hNOE≈0.9) upon ET binding and closure of the cleft (Fig. 3, fig. S9) (*16*). This is a result of R379 NHε donating an H-bond to D381 Oδ1 (Fig. 3D, 3E) (*43*). Strikingly, although the chemical shift of this crosspeak in the I243P mutant is WT-like, the intensity is far weaker (Fig. 3E) and the ^15^N-^1^H-hNOE measurably smaller (fig. S9). These data reveal a weaker H-bond in the mutant in the context of the WT-like binding cleft (*44*).

### Molecular dynamics simulations of WT vs. I243P *Sp*EgtUC show dynamical changes upon perturbation of CH•••S H-bonds

The I243P *Sp*EgtUC mutant exhibits a wild-type-like ET binding pocket with at least one weaker H-bond and a dramatically shorter “closed”-state lifetime relative to the WT domain. To address what structural changes in the binding pocket result in the shorter “closed”-state lifetime, we carried out 1 µs all-atom molecular dynamics (MD) simulations of the solvated WT (movie S1) and I243P *Sp*EgtUC (movie S2) ET-bound complexes to obtain a time-course view of the ligand dynamics at the binding site in each complex. We parameterized ET consistently with the OPLS-AA protein force field, which showed a C-S double bond (1.629Å) in the thioimidazole ring and a significant partial negative charge (–0.49) on the S atom. Both Nδ1 and Nε2 are fully protonated in this thione conformer as anticipated (Fig. 1C).

Interestingly, while the WT-ET complex remains closed during the 1 µs simulation (Fig. 4A, complex A (movie S1), the I243P mutant complex dissociates, forming a pre-dissociation state after 660 ns (Fig. 4A, complex B), followed by full ET dissociation after 860 ns (Fig. 4A, complex C) (movie S2). Thus, the MD simulation of the I243P mutant complex captures a rare ET dissociation event. The pre-dissociation state of the transporter is characterized by a more “open” ligand binding site due to a movement of the D2 lobe relative to D1; D2 harbors E375, R379 and D381 that form an H-bond network on one side of the ligand.

**Figure 4.**
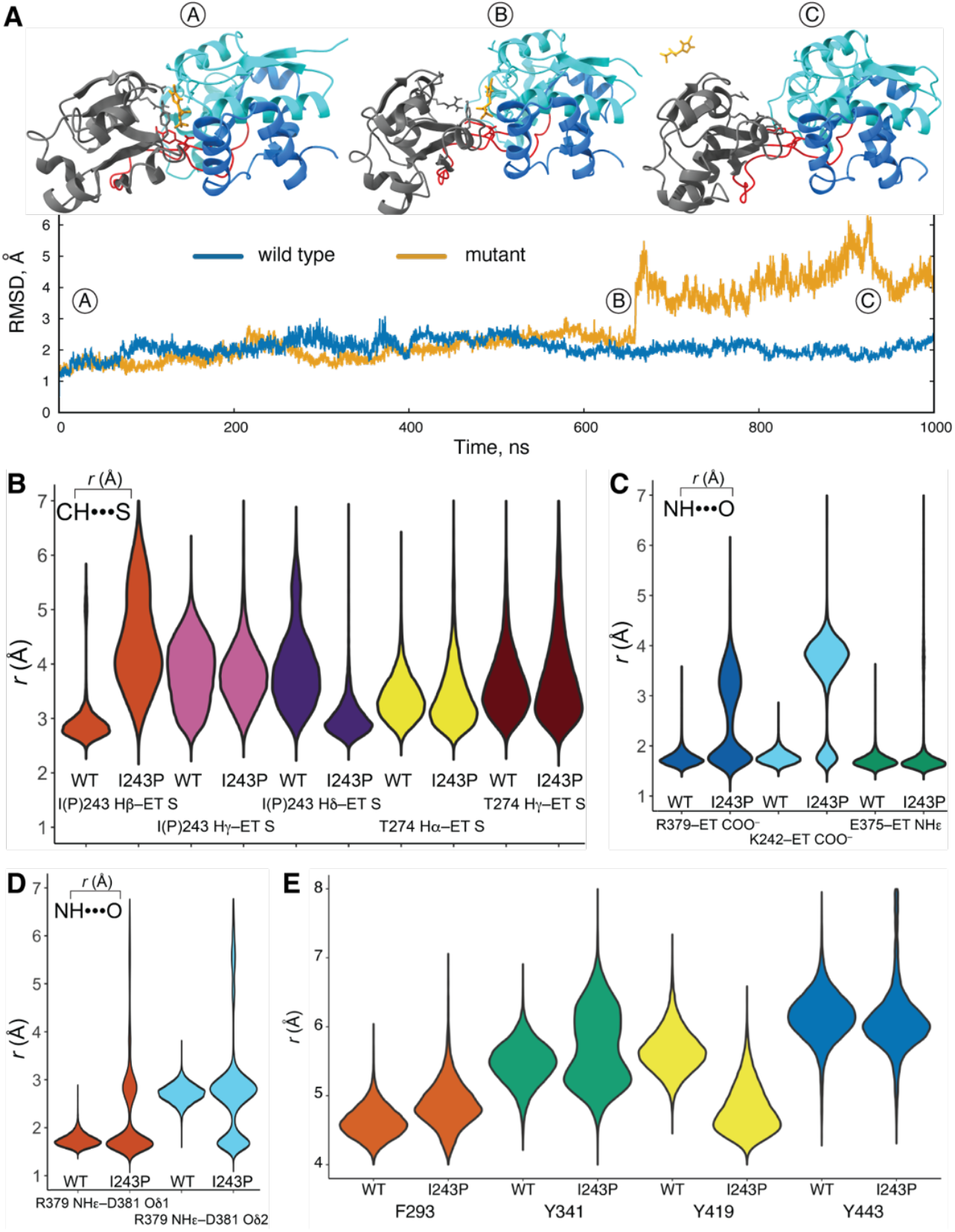
Molecular dynamics simulations of WT and I243P *Sp*EgtUC. (**A**) RMSD (Å) of WT (blue) (movie S1) and I243P (yellow) (movie S2) *Sp*EgtUC over a 1 µs simulation representing [A] the closed state, [B] the pre-dissociation state and [C] the fully dissociated state. (**B-E**) Violin (histogram) plots of closed-state bond distances for WT EgtUC and I243P EgtUC. (**B**) C-H•••S hydrogen bond distances for the hydrogens of residue 243 (Hβ, orange, Hɣ, pink, and Hδ, purple), or T274 (H⍺, yellow, and Hɣ, brown). (**C**) N-H•••O hydrogen bond distances for R379 and COO^-^ of ET (blue), K242 and COO^-^ of ET (cyan), and E375 and NHε of ET (green). (**D**) N-H•••O hydrogen bond distances for the NHε of R379 and D381 Oδ1 (orange) or D381 Oδ2 (cyan). (**E**) Cation-π distances from the trimethylamine moiety of ET to the center of the aromatic cage residues F293 (orange), Y341 (green), Y419 (yellow), and Y443 (blue).

We next carried out a time-course analysis of H-bond length distributions, focusing on the “closed” state of each bound complex (0-1 µs for WT; 0-660 ns for I243P EgtUC). The distance distributions of candidate alkyl CH•••S H-bonds in the WT domain reveals that the I243 Hβ-ET distance is characterized by very rare long-length deviations, fluctuating close to the average value of ≈2.8 Å (Fig. 4B) over the course of the simulation. Other CH•••S H-bonds behave similarly, including the T274 H⍺-ET and T274 Hɣ-ET interactions (Fig. 4B). Remarkably, the same holds true for the closed form of the I243P mutant complex where the primary interaction of the S atom is with the P243 Hδ hydrogen atom (Fig. 4B). We conclude that the I243P mutant CH•••S H-bond dynamics parallel the WT complex, suggesting that these CH•••S H-bonds are stable and contribute comparably to the interaction of ET with the EgtUC binding pocket in both domains.

A corresponding analysis of the conventional NH•••O H-bonding interactions reveals that the R379-ET COO^−^ and K242-ET COO^−^ H-bond distances in the WT complex exhibit narrow single-peak distributions (Fig. 4C) and thus appear stable during the simulation. In striking contrast, the same H-bond distance distributions in the I243P mutant are bimodal, demonstrating that these H-bonds are effectively broken for a significant fraction of the simulation time, portending the pre-dissociation state (Fig. 4C; movie S2). A notable exception to this behavior is the ET Nε2-H•••E375 Oε1 H-bond whose distance distributions appear similar in the WT and I243P domains (Fig. 4C). In contrast, the ET carboxylate moiety experiences considerable disorder in the I243P mutant complex, implicating a more disordered H-bonding network in the I243P binding pocket. In WT EgtUC, indeed, R379 NHε donates a H-bond exclusively to D381 Oδ1 (1.8 Å), with Oδ2 consistently further away (2.8 Å) (Fig. 4D). Strikingly, in I243P *Sp*EgtUC, both distances are significantly populated, suggesting rotation of the carboxylate side chain of D381 in the mutant (Fig. 4D). The simulated dynamics are fully consistent with the NMR data that suggest that the R379 NHε-D381 Oδ1 interaction is weakened in the I243P mutant (Fig. 3E; fig. S9).

The cation-π interactions between the trimethylamine moiety and the aromatic cage at the base of the binding site, like the CH•••S distance distributions, are largely unperturbed in the mutant vs. wild-type simulations (Fig. 4E). While the interaction between ET and Y341 shows a bimodal distribution in the mutant, the other residues are either unperturbed or appear to show compensatory behavior. We conclude that strong alkyl CH•••S H-bonds anchor the ET-EgtUC complex, and nucleate conventional electrostatic and cation-π interactions distant from the thioimidazole ring. Even small perturbations in these CH•••S interactions dramatically impact the complex lifetime due largely to weakening of the ET carboxylate “clamp”. This explains why HER, which lacks the thione S altogether (fig. S2A) binds so weakly to EgtUC (≥10-fold weaker than ET binds to the I243P mutant) (*16*) despite the full complement of these other H-bonding interactions.

### EgtUC mutants exhibit reduced ET uptake in *S. pneumoniae* cells

We next tested if an intact EgtU transporter that harbors an I243 EgtUBC substitution, relative to WT and WT-like T274G mutant, might be functionally compromised in its ability to uptake ET into cells. To do this, we constructed T274G, I243A and I243P *egtUBC* mutant strains of *S. pneumoniae* D39 via allelic exchange into the native chromosomal locus. We first grew these cells on a chemically defined (C) media (*45*) supplemented with variable concentrations of ET and used a single endpoint assay (3 h) to assess the impact of these mutations on ET accumulation in *S. pneumoniae* D39 using a quantitative isotope-dilution thiol profiling approach (Fig. 5A) (*16*). All four strains behave similarly in this assay, with ET accumulation slightly less efficient in the I243P mutant at the lowest concentration of ET tested (50 nM), far below the *K*_d_ (Table 1, Fig. 5A). To address if this small difference in total ET at 3 h might be more dramatic at earlier timepoints, we investigated the kinetics of ET uptake by the WT and I243P strains, following the addition (*t*=0) of a low, yet physiologically relevant concentration of ET (50 nM). We find that in WT *S. pneumoniae*, ET accumulates to high intracellular concentrations and is fully depleted from the media in ≈10 min, as indicated by decreasing intracellular ET at later timepoints (Fig. 5B); a similar finding was made in *L. reuteri* (*28*). Strikingly, the I243P *egtUBC* strain accumulates ET to significantly lower extents at all time points (Fig. 5B; fig. S10). These biological findings reveal that while the mutant transporter remains functional, reduced lifetime of transport-competent “closed” state in I243P *Sp*EgtUC (Fig. 3-4) negatively impacts cellular ET uptake.

**Figure 5.**
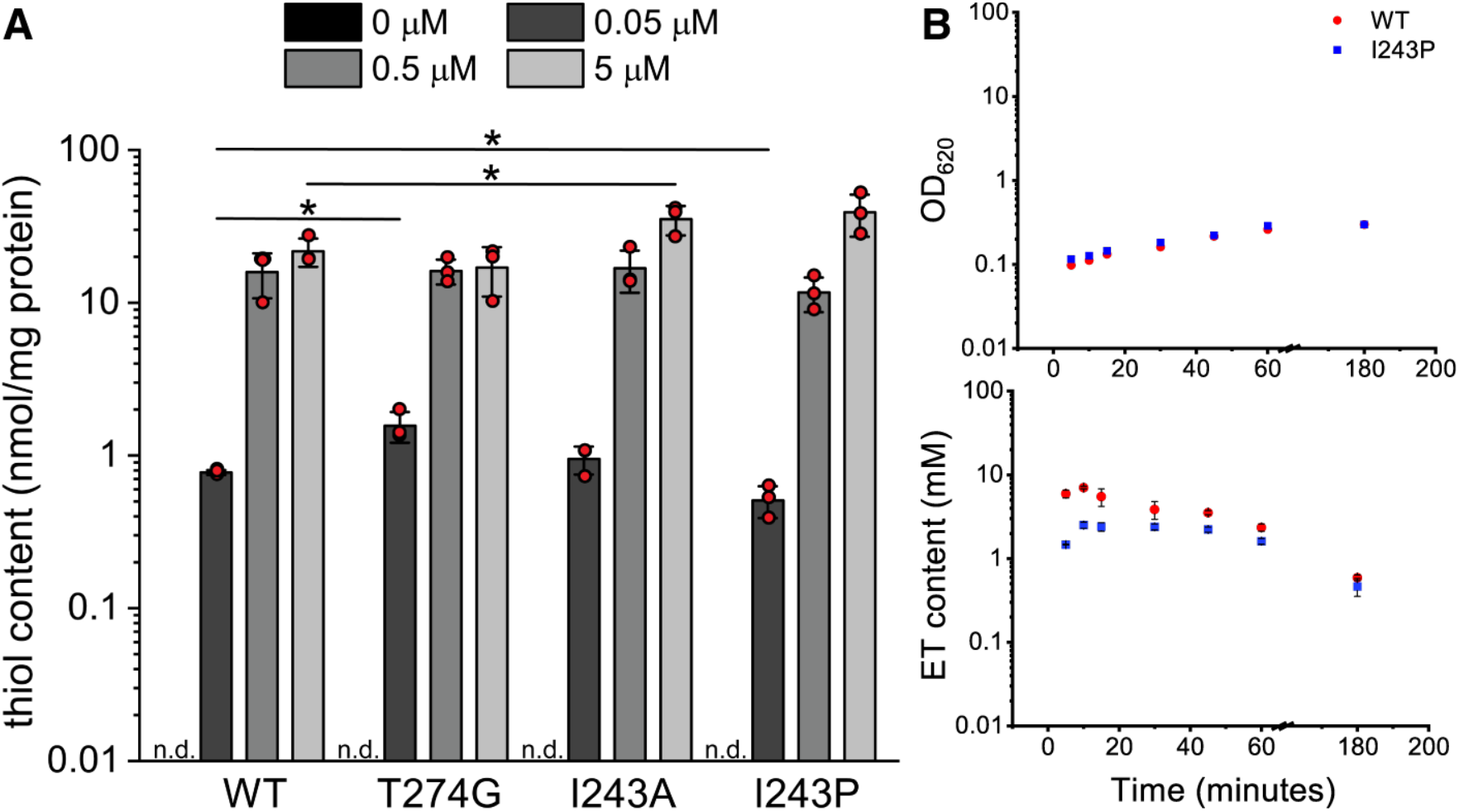
LMW-thiol profiling of EGT in *S. pneumoniae* strains. *Left to right:* Ergothioneine profiling of wild-type, T274G, I243A, and I243P EgtUBC mutants grown in chemically defined media (CDM) (*45*) with increasing concentrations of extracellular ET (0, 0.05, 0.5, 5 µM) taken a 3-h growth time point. ^*^*p*<0.05 (**B**) The OD_620_ (*upper*) and kinetics of ET uptake (l*ower*) for the wild-type (red circles) and I243P (blue squares) *egtUBC* strains following the addition of 0.05 µM ET at *t*=0, with the results of three technical replicates shown (see also fig. S10). The 180 min data are taken from panel (A).

## Discussion

In this work, we exploit molecular recognition of ET by *Sp*EgtUC to show that a limited set of alkyl CH•••S hydrogen bonds dramatically impact metabolite recognition by a protein host in aqueous solution. The contributions of H-bonding to ET binding affinity, donated by both the backbone and the side chains, reveal that Hα, methine and methylene hydrogens are stronger H-bond donors to the S atom than are methyl hydrogens, which tracks with the polarizability of the C-H bond (*36*). A particularly striking example is the wild-type ET binding affinity of the T274G mutant, where despite the fact that both the γ-methyl hydrogens and the Hα are in close physical proximity to the S atom, each meeting the empirical structural criteria for a strong CH•••S H-bond (fig. S5B, middle) (*36*), it is the backbone Hα of T274 that makes the energetically important H-bond. The I243A substitution reinforces this trend, with the methyl group donor in I243A EgtUC destabilized by about ≈2.0 kcal mol^-1^, relative the β-methine hydrogen of WT I243, despite possessing the nearly identical H•••S distance and CH•••S bond angle characteristics (Table 1). The affinity of the I243A mutant (but not the underlying energetics) is comparable to that of the distantly related *Hp*EgtUC which harbors a Pro in this position. The behavior of the I243P mutant in the *Sp*EgtUC series also reinforces this general trend in H-bond strength but exhibits a more complex behavior. The complexity arises because one of the Hδ hydrogens of P243 makes closest approach to the S atom in these two cases, while it is the Hβ hydrogen of I243 that performs this role in the WT domain. In addition, when the I243P mutant structure is analyzed in more detail, we find increased CH•••S hydrogen bond lengths and slightly more acute bond angles (Table 2), consistent with weaker H-bonding interactions (*36*). Indeed, the Pro substitution is strongly destabilizing, far more so than in *Hp*EgtUC itself, and we were unable to recover *Hp*EgtUC binding affinity in the double I243P/T274G “*Hp*-like” *Sp*EgtUC mutant (Table 1).

Remarkably, our structures collectively reveal only small perturbations in the binding pocket, despite *K*_a_ values that vary over three orders of magnitude. These small perturbations manifest as significantly reduced enthalpies of binding, shorter closed-state lifetimes, and increased time-averaged disorder in the I243P vs. WT domains. Remarkably, this dynamical disorder is centered not on the CH•••S H-bonding partners, but on the conventional NH•••O H-bonds that stabilize the electrostatic “clamp” around the ET carboxylate moiety. Indeed, a predissociation state observed only in the I243P mutant simulations reveals that these weakened interactions are lost altogether, which after some time, all thioimidazole ring interactions are lost, resulting in complete dissociation of the ligand. These perturbations negatively impact the kinetics of ET uptake into *S. pneumoniae*, a finding that has strong implications for competition for this dietary antioxidant in a mixed microbial community or in competition with host cells (*28*). Indeed, *H. pylori* EgtU transports ET efficiently, despite possessing an ≈40-fold lower reported affinity than *Sp*EgtUC (Table 1) (*17*).

The nucleating CH•••S H-bonds described here ensure high ET specificity by EgtU, which appears to contrast with the known characteristics of the ET-transporting organic cation transporter novel 1 (OCTN1), which is generally considered a non-selective carrier of organic cations, anions and drug molecules (*46*). Additional insights from ongoing biochemical, mechanistic and structural studies of intact EgtU promises to provide new insights into microbial ET transport, while potentially enabling the development of a broad spectrum EgtU-selective inhibitor as a novel antibiotic strategy against human microbial pathogens.

## Acknowledgements

The authors gratefully acknowledge use of the Macromolecular Crystallography Facility (MCF) and the Advanced Research Computing Facilities at Indiana University Bloomington. We also thank Jay Nix for his assistance during X-ray data collection at beamline 8.2.2 at Advanced Light Source (ALS), Berkeley, CA. The 800 MHz NMR spectrometer used in this research in the METACyt Biomolecular NMR facility at Indiana University Bloomington was supported in part by a grant from the NIH (S10 OD032431-01A1). We thank Prof. Malcolm Winkler and Dr. H.C. Tiffany Tsui (Indiana University, Bloomington) for resources supported by a grant from NIGMS (R35 GM131767).

## Author contributions

Conceptualization: DPG, KAL

Formal analysis: KAL

Funding acquisition: DPG

Investigation: KAL, GG-G, KAE, PGS.

Supervision: DPG.

Validation: DPG, KAL, KAE.

Visualization: DPG, KAL, KAE, PGS

Writing- original draft: DPG, KAL, KAE, PGS

Writing- review and editing: DPG, KAL, KAE, PGS, GG-G

## Data and materials availability

Raw data and materials described in this work are available upon request from the authors.

## Supplementary Materials

Materials and Methods

Figs. S1 to S9

Tables S1 to S3

Movies S1 to S

## References

1. R. C. Fahey, Glutathione analogs in prokaryotes. Biochim Biophys Acta 1830, 3182–3198 (2013).

2. A. Gaballa et al., Biosynthesis and functions of bacillithiol, a major low-molecular-weight thiol in Bacilli. Proc Natl Acad Sci U S A 107, 6482–6486 (2010).

3. A. M. Reyes et al., Chemistry and redox biology of mycothiol. Antioxid Redox Signal 28, 487–504 (2018).

4. K. Ulrich, U. Jakob, The role of thiols in antioxidant systems. Free Radic Biol Med 140, 14–27 (2019).

5. M. L. Reniere, Reduce, induce, thrive: Bacterial redox sensing during pathogenesis. J Bacteriol 200, e00128–00118 (2018).

6. A. Anaya-Sanchez, Y. Feng, J. C. Berude, D. A. Portnoy, Detoxification of methylglyoxal by the glyoxalase system is required for glutathione availability and virulence activation in Listeria monocytogenes. PLoS Pathog 17, e1009819 (2021).

7. Z. Rosario-Cruz, J. M. Boyd, Physiological roles of bacillithiol in intracellular metal processing. Curr Genet 62, 59–65 (2016).

8. D. G. Dumitrescu, S. K. Hatzios, Emerging roles of low-molecular-weight thiols at the host-microbe interface. Curr Opin Chem Biol 75, 102322 (2023).

9. J. M. Lensmire et al., The glutathione import system satisfies the Staphylococcus aureus nutrient sulfur requirement and promotes interspecies competition. PLoS Genet 19, e1010834 (2023).

10. P. J. Kies, N. D. Hammer, A resourceful race: Bacterial scavenging of host sulfur metabolism during colonization. Infect Immun 90, e0057921 (2022).

11. T. T. Fu, L. Shen, Ergothioneine as a natural antioxidant against oxidative stress-related diseases. Front Pharmacol 13, 850813 (2022).

12. Y. Gao et al., L-Ergothioneine exhibits protective effects against dextran sulfate sodium-induced colitis in mice. ACS Omega 7, 21554–21565 (2022).

13. B. Halliwell, I. K. Cheah, R. M. Y. Tang, Ergothioneine - a diet-derived antioxidant with therapeutic potential. FEBS Lett 592, 3357–3366 (2018).

14. C. M. Kayrouz, J. Huang, N. Hauser, M. R. Seyedsayamdost, Biosynthesis of selenium-containing small molecules in diverse microorganisms. Nature 610, 199–204 (2022).

15. K. A. Edmonds, K. Diaz-Rodriguez, D. P. Giedroc, Expression and purification of the intact bacterial ergothioneine transporter EgtU. Protein Expr Purif 227, 106633 (2025).

16. Y. Zhang et al., Discovery and structure of a widespread bacterial ABC transporter specific for ergothioneine. Nat Commun 13, 7586 (2022).

17. D. G. Dumitrescu et al., A microbial transporter of the dietary antioxidant ergothioneine. Cell 185, 4526-4540.e4518 (2022).

18. G. Barger, A. J. Ewins, The constitution of ergothioneine, a betaine related to histidine. J Am Chem Soc 99, 2336–2341 (1911).

19. D. Grundemann, L. Hartmann, S. Flogel, The ergothioneine transporter (ETT): substrates and locations, an inventory. FEBS Lett, (2021).

20. I. Tamai et al., Cloning and characterization of a novel human pH-dependent organic cation transporter, OCTN1. FEBS Lett 419, 107–111 (1997).

21. D. Grundemann et al., Discovery of the ergothioneine transporter. Proc Natl Acad Sci U S A 102, 5256–5261 (2005).

22. B. M. Cumming, K. C. Chinta, V. P. Reddy, A. J. C. Steyn, Role of ergothioneine in microbial physiology and pathogenesis. Antioxid Redox Signal 28, 431–444 (2018).

23. K. A. Jenny, G. Mose, D. J. Haupt, R. J. Hondal, Oxidized forms of ergothioneine are substrates for mammalian thioredoxin reductase. Antioxidants (Basel) 11, 185 (2022).

24. L. Hartmann, F. P. Seebeck, H. G. Schmalz, D. Grundemann, Isotope-labeled ergothioneine clarifies the mechanism of reaction with singlet oxygen. Free Radic Biol Med 198, 12–26 (2023).

25. D. Petrovic et al., Ergothioneine improves healthspan of aged animals by enhancing cGPDH activity through CSE-dependent persulfidation. Cell Metab 37, 542-556.e514 (2025).

26. M. Richard-Greenblatt et al., Regulation of ergothioneine biosynthesis and its effect on Mycobacterium tuberculosis growth and infectivity. J Biol Chem 290, 23064–23076 (2015).

27. V. Saini et al., Ergothioneine maintains redox and bioenergetic homeostasis essential for drug susceptibility and virulence of Mycobacterium tuberculosis. Cell Rep 14, 572–585 (2016).

28. I. K. Cheah, J. Z. Lee, R. M. Y. Tang, P. W. Koh, B. Halliwell, Does Lactobacillus reuteri influence ergothioneine levels in the human body? FEBS Lett 596, 1241–1251 (2022).

29. M. A. Beliaeva, F. Leisinger, F. P. Seebeck, In vitro reconstitution of a five-step pathway for bacterial ergothioneine catabolism. ACS Chem Biol 16, 397–403 (2021).

30. H. Muramatsu et al., Characterization of ergothionase from Burkholderia sp. HME13 and its application to enzymatic quantification of ergothioneine. Appl Microbiol Biotechnol 97, 5389–5400 (2013).

31. Z. Zhou, S. K. Hatzios, Microbial metabolism of host-derived antioxidants. Curr Opin Chem Biol 84, 102565 (2025).

32. B. C. Chu, T. DeWolf, H. J. Vogel, Role of the two structural domains from the periplasmic Escherichia coli histidine-binding protein HisJ. J Biol Chem 288, 31409–31422 (2013).

33. I. Smirnova, V. Kasho, H. R. Kaback, Real-time conformational changes in LacY. Proc Natl Acad Sci U S A 111, 8440–8445 (2014).

34. S. J. Ruiz, G. K. Schuurman-Wolters, B. Poolman, Crystal structure of the substrate-binding domain from Listeria monocytogenes bile-resistance determinant BilE. Crystals 6, 162 (2016).

35. I. K. Cheah, R. M. Tang, T. S. Yew, K. H. Lim, B. Halliwell, Administration of Pure Ergothioneine to healthy human subjects: uptake, metabolism, and effects on biomarkers of oxidative damage and inflammation. Antioxid Redox Signal 26, 193–206 (2017).

36. H. A. Fargher, T. J. Sherbow, M. M. Haley, D. W. Johnson, M. D. Pluth, C-H•••S hydrogen bonding interactions. Chem Soc Rev 51, 1454–1469 (2022).

37. H. A. Fargher et al., Tuning supramolecular selectivity for hydrosulfide: Linear free energy relationships reveal preferential C-H hydrogen bond interactions. J Am Chem Soc 142, 8243–8251 (2020).

38. A. G. Davis, L. N. Zakharov, M. D. Pluth, Probing the reversible binding of anionic reactive sulfur and nitrogen species in imidazolium receptors with directional C-H hydrogen bonds. Inorg Chem 64, 7774–7783 (2025).

39. A. G. Davis, L. N. Zakharov, M. D. Pluth, Reversible hydrosulfide (HS(-)) ninding using exclusively C-H hydrogen-bonding interactions in imidazolium hosts. Inorg Chem 63, 3057–3062 (2024).

40. M. W. Krone et al., Thermodynamic consequences of Tyr to Trp mutations in the cation-pi-mediated binding of trimethyllysine by the HP1 chromodomain. Chem Sci 11, 3495–3500 (2020).

41. A. Paleskava, A. L. Konevega, M. V. Rodnina, Thermodynamics of the GTP-GDP-operated conformational switch of selenocysteine-specific translation factor SelB. J Biol Chem 287, 27906–27912 (2012).

42. M. K. Osterberg et al., Coupling of zinc and GTP binding drives G-domain folding in Acinetobacter baumannii ZigA. Biophys J 123, 979–991 (2024).

43. D. Nguyen, C. Chen, B. M. Pettitt, J. Iwahara, NMR methods for characterizing the basic side chains of proteins: Electrostatic interactions, hydrogen bonds, and conformational dynamics. Methods Enzymol 615, 285–332 (2019).

44. M. A. Pennella, A. I. Arunkumar, D. P. Giedroc, Individual metal ligands play distinct functional roles in the zinc sensor Staphylococcus aureus CzrA. J Mol Biol 356, 1124–1136 (2006).

45. I. van de Rijn, R. E. Kessler, Growth characteristics of group A streptococci in a new chemically defined medium. Infect Immun 27, 444–448 (1980).

46. O. Ben Mariem et al., Atomistic description of the OCTN1 recognition mechanism via in silico methods. PLoS One 19, e0304512 (2024).

47. T. Lundback, S. van Den Berg, T. Hard, Sequence-specific DNA binding by the glucocorticoid receptor DNA-binding domain is linked to a salt-dependent histidine protonation. Biochemistry 39, 8909–8916 (2000).

48. D. Sehnal et al., Mol* Viewer: modern web app for 3D visualization and analysis of large biomolecular structures. Nucleic Acids Res 49, W431–W437 (2021).

