## Supplemental Materials for "CH•••S hydrogen bonds drive molecular recognition of ergothioneine by the microbial transporter"

**The PDF file includes:**

Materials and Methods
Figs. S1 to S10
Tables S1 to S5
Captions for Movies S1 to S2

**Other Supplementary Materials for this manuscript include the following:**

Movies S1 to S2

### Materials and Methods

#### Materials and chemicals

*L*-glutathione (GSH) was obtained from Sigma Aldrich (G4251), *L*-ergothioneine was obtained from Santa Cruz Biotechnology (sc-200814), *L*-cysteine from Fisher Biotech (BP376-100), *L*-hercynine from Toronto Research Chemicals (H288900), L-histidine from Sigma Aldrich (H8000), *β*-(4-hydroxyphenyl) ethyl iodoacetamide from Chem-Impex, and *N*-iodoacetyltyramine-*d*4 from Toronto Research Chemicals (I685882). These chemicals were used without further purification. Other chemicals include IPTG from GoldBio (2481C100), Imidazole from Chem-Impex (00418), dithiothreitol from Chem-Impex (00127), HEPES from Chem-Impex (00174), Tris-HCl from MP Biomedicals (816100), EDTA from VWR Chemicals (BDH9232), and NaCl from VWR Chemicals (BDH9286). BHI broth was obtained from BD (37500) while Luria broth was obtained from Fisher Bioreagent (BP9723). Milli-Q water was used to make all solutions.

#### Gene block sequence for I243P/T274G EgtUC double mutant

GATCCCATATGGAGAAGGAAAACTTGATTATTGCTGGGAAACCGGGCCCAGA
ACCAGAAATTTTGGCTAATATGTATAAATTGCTGATTGAAGAAAATACCAGCATGAC
TGCGACTGTTAAACCGAATTTTGGGGGGACAAGTTTCCTTTATGAAGCTCTGAAAAA
AGGTGATATTGACATTTATCCTGAATTTACCGGTACGGTGACTGAAAGTTTGCTTCA
ACCATCACCCAAGGTGAGTCATGAGCCAGAGCAGGTTT

#### Mutagenesis, protein expression, and protein purification

Mutagenesis, expression, and purification of *Sp*EgtUC mutants was performed as described previously for wild-type *Sp*EgtUC using the mutagenic primers listed in table S4 (*1*). Briefly, transformed cells were grown at 37°C with shaking until $OD_{600}$=0.8. IPTG was added to a final concentration of 1 mM. Cells were then grown at 18 °C for 18 h. Cell pellets were resuspended in 40 mL Buffer A (25 mM Tris pH 8.0, 20 mM imidazole, 500 mM NaCl, 10% glycerol) and lysed by sonication using a sonic dismembrator, 65% amplitude, 2.0 s on, 8.0 s off, for 5 minutes total lysis time. The cell lysate was clarified by centrifugation and ammonium sulfate was added to the supernatant to a final concentration (w/v) of 70%. The ammonium sulfate pellet was resuspended in 35 mL Buffer A and loaded onto a 5 mL Ni-NTA affinity column at a flow rate of 5 mL/min, controlled by an AKTA Pure chromatography system. EgtUC was eluted using an imidazole gradient (0-100% B). Fractions were collected and dialyzed overnight in 1 L buffer A supplemented with 2 mM DTT. SUMO protease was added to fractions to cleave the $His_6$-SUMO tag. The following day, the dialysate was again loaded onto Ni-NTA column and the flow-through fractions collected. These fractions were pooled and further chromatographed by size exclusion using a HiLoad 16/600 Superdex 75 column at a flow rate of 1 mL/min. EgtUC-containing fractions as assessed by SDS-PAGE were pooled conservatively and used without further purification. Pooled samples were subjected to ESI-MS to confirm protein identity. Expected and observed masses for all recombinant *Sp*EgtUC proteins used in this study are listed below (mass accuracy ±5 Da).

| Protein | Expected mass (Da) | Observed mass (Da) |
|---|---|---|
| WT *Sp*EgtUC | 31032.41 | 31027 |
| I243A | 30990.33 | 30985 |

| | | |
|---|---|---|
| I243P | 31016.37 | 31016 |
| T274G | 30988.36 | 30983 |
| K242A | 30975.31 | 30970 |
| K242S | 30991.31 | 30986 |
| I243P/T274G | 30972.31 | 30972 |

**Isothermal titration calorimetry**

ITC experiments were carried out using a MicroCal VP-ITC calorimeter at 25 (±0.1) °C by titrating 20, 30, or 200 μM wild-type or mutant *Sp*EgtUC in the sample chamber in 50 mM HEPES, pH 7.5, 150 mM NaCl, 2 mM EDTA with the indicated ligand (ET or *L*-hercynine) dissolved in the syringe in same buffer. For the ET titration, the ligand concentration in the syringe was typically 375 μM with 30 μM protein in the sample chamber, or 2.25 mM with 200 μM protein in the sample chamber for I243P and I243P/T274G EgtUC. For hercynine, the ligand concentration was 600 μM, with 20 μM protein in the sample chamber. For temperature dependence studies, experiments were carried out at 15, 20, 30, and 35 °C using 10 μM WT EgtUC and 100 μM ET. The data analysis package provided by MicroCal was utilized to integrate raw ITC data and plot as heat versus ligand to protein ratio in Origin. Signals for heats of dilution obtained for titration of ligand into buffer were subtracted from raw data prior to analysis. Corrected data were fit to a single site binding model using the data analysis package, allowing *n*, the number of binding sites, to float.

**Temperature-dependence of ET binding calculations**

Temperature-dependence of ET binding was calculated as described previously (*2*). Briefly, we performed an unweighted linear fit to the enthalpy vs. temperature plot (Fig. 1E, bottom) to obtain a value for $\Delta C_p = -0.29$ kcal mol$^{-1}$ K$^{-1}$ for binding of *Sp*EgtUC to ET (assuming a temperature-independent change in heat capacity), and a temperature ($T_H$) for which $\Delta H(T_H)=0$; $T_H = 263.1$ K. We then fit $\Delta G°$ as a function of temperature to eq 1:

$$\Delta G° = \Delta C°_p[T\left(1 - ln\frac{T}{T_s}\right) - T_H \quad (1)$$

where $\Delta C_p°$ and $T_H$ were fixed to the values indicated with a single adjustable parameter, $T_s$ when $\Delta S(T_S)=0$. We determined $T_s = 298.9$ K, which corresponds to the temperature at which ΔG° is at a minimum (most negative) value.

**Solvent accessible surface area calculations**

The change in solvent accessible surface area of *Sp*EgtUC binding to ET was calculated using a published method. Briefly, the change in constant pressure heat capacity is broken down into two parts (eq 2):

$$\Delta C°_p = \Delta C_{ap}\Delta SASA_{ap} + \Delta C_p \Delta SASA_p \quad (2)$$

where $\Delta C_{ap}$ and $\Delta C_p$ are the apolar and polar contributions, respectively, to the heat capacity of the system. Using previously established values for $\Delta C_p$ and $\Delta C_{ap}$ (*3*) we estimated the change in solvent accessible surface area (ΔSASA) assuming all apolar or all polar residues, and using the average size of apolar and polar residues, calculated the number of amino acids buried during binding of *Sp*EgtUC to ET (Table 3).

**X-ray crystallography**

T274G, I243A, I243P and I243P/T274G EgtUC in complex with ET at ~15 mg/mL were crystallized following the procedure published before (*1*). Crystals grew in tribasic sodium citrate, pH 5.6, 1.2 – 1.8 M at 20°C using the sitting-drop vapor-diffusion method. Crystals were harvested, washed for few seconds in reservoir solution and flash-frozen in liquid nitrogen. Diffraction data were collected at 100 K at the Beamline station 8.2.2 at the Advanced Light Source (Berkeley National Laboratory, CA) and were initially indexed, integrated, and scaled using XDS (*4*). Molecular replacement was used to estimate phases using PHASER and PDB code 7TXK(*1*) as search model. Data sets for mutants I243A EgtUC and T274G EgtUC presented moderate to severe anisotropy, with useful data extending to ≈2 Å (see table S1). Thus, data sets were re-processed, and anisotropy analysis was performed using the STARANISO server (*5*). Anisotropic completeness was obtained by fitting an ellipsoid using least squares to the resulting cut-off surface points. Cut-off criteria were Rpim < 0.6, I/s > 1.2 and CC1/2 > 0.5. Final data sets, after anisotropic correction, were used to perform molecular replacement. Successive cycles of automatic building in Autobuild (PHENIX) and manual building in Coot, as well as refinement (PHENIX Refine) led to complete models (*6, 7*). MolProbity software (*8*) was used to assess the geometric quality of the models, and Pymol was used to generate molecular images. Data collection and refinement statistics are indicated table S1.

**NMR spectroscopy**

Uniformly $^{15}N$-labeled protein was expressed in *E. coli* BL21 (DE3) cells in M9 medium containing 1.0 g/L of $^{15}NH_4Cl$ as the sole nitrogen source. Expression, isolation and purification of these isotope-labeled proteins was performed as described above for unlabeled protein. $^{15}N$ TROSY spectra of WT EgtUC were recorded 35 °C on an 800 MHz Bruker Avance Neo spectrometer equipped with a cryogenic probe in the METACyt Biomolecular NMR Laboratory at Indiana University. Titration of WT EgtUC with ET was performed with 0.2 mM protein in 10 mM sodium phosphate pH 7.0, 150 mM NaCl, and 10% v/v $D_2O$, with 0.3 mM 2,2-dimethyl-2-silapentanesulfonic acid (DSS) as an internal reference. Final concentrations of ET were 0, 50, 100, 150, 200, and 250 µM ET. $^{15}N$ TROSY spectra of EgtUC mutants were recorded at 35 °C on a 600 MHz Bruker Avance Neo spectrometer equipped with a cryogenic probe. Titration of I234P EgtUC with ET was performed with 0.4 mM protein in 10 mM sodium phosphate pH 7.0, 150 mM NaCl, and 10% v/v $D_2O$, with 0.3 mM 2,2-dimethyl-2-silapentanesulfonic acid (DSS) as an internal reference. Final concentrations of ET were 0, 0.0125, 0.025, 0.05, 0.1, 0.2, 0.4, 0.8, 1.6, 3.2, 6.4 and 12.8 mM ET. A titration of I234A EgtUC was performed with 0.1 mM protein in the same buffer, with final concentrations of 2, 4, 8, 16, 25, 50, 75, 100, 125, 150, 175, 200, 250, and 300 µM ET. The $^{15}N$ spin relaxation rate $R_2$ and $^{1}H$-$^{15}N$ heteronuclear NOE (hNOE) values were measured using TROSY pulse sequences as described previously.(*1*) The relaxation delays used were 0.017, 0.034, 0.051, 0.068, 0.085, 0.102, 0.119, 0.136, 0.170, and 0.204 s for $R_2$. Residue-specific $R_2$ values were obtained from fits of peak intensities vs. relaxation time to a single exponential decay function, while hNOE ratios were ascertained directly from intensities in experiments recorded with (2 s relaxation delay followed by 3 s saturation) and without saturation (relaxation delay of 5 s). Errors in hNOE values were calculated by propagating the error from the signal to noise ratio.

**Molecular dynamics simulations**

We modeled the protein using the OPLS-AA force field (*9*) and the explicit water molecules using the SPC/E model (*10*). We derived parameters for ET consistent with the OPLS-

AA force field using LigParGen (*11*). We employed the crystallographic coordinates as starting configurations for the WT and I243P mutant ligand-protein complexes together with resolved crystallographic water molecules, which we then solvated in a cubic box of explicit water molecules centered around the protein-ligand complex, allowing for at least 1 nm from the box edges. We neutralized the simulation system using $Na^+$ ions, which we placed at random positions in the box. We first optimized the starting geometries of each system and then equilibrated the simulation box at temperature 300 K and pressure 1 bar by running a 100 ps NVT simulation followed by 100 ps NPT simulation, including position restraints on the heavy atoms of the protein-ligand complex. Following equilibration, we switched off the position restraints and ran the productive MD simulations for 1 µs with a sampling step size of 40 ps, resulting in 25,000 snapshot MD trajectories. For all MD runs, we used GROMACS (*12-14*) with a 2 fs timestep, constrained the hydrogen-containing covalent bonds using LINCS (*15, 16*) and used 1 nm short-range cut-off for non-covalent interactions and accounted for long-range electrostatic interactions using PME (*17*), applied the modified Berendsen thermostat (*18*) at a reference temperature of 300 K, a time constant of 0.1 ps and two coupling groups for the protein-ligand complex and the solvent, applied the Parrinello-Rahman barostat (*19*) at a reference pressure of 1 bar, a time constant of 2 ps and isotropic rescaling. We generated movies of the trajectories using ChimeraX (*20*) by centering and fitting the protein-ligand configurations to the initial configuration using gmx trjconv to remove translational and rotational diffusion. We computed inter-atomic distances using gmx distance in GROMACS and generated violin plots using ggplot2 (*21*). We also computed averaged per residue solvent-accessible surface-areas of the WT protein-ligand complex using gmx sasa (*22*) with a probe radius of 0.14 nm, and compared the results with the averaged per residue solvent-accessible surface-areas of the apo protein, which we equilibrated using the same protocol as above and simulated for 100 ns.

#### ***Streptococcus pneumoniae*** **D39 mutant strain construction and LMW thiol quantification in cells**

Mutant strains listed in table S3 were constructed using standard laboratory practices for allelic replacement in *S. pneumoniae* serotype 2 D39W (IU1781) (*1*). All mutant strain constructs were sequence verified. Primers are listed in table S4-S5. Strains were grown overnight in a chemically defined media (CDM) lacking ET (*23*). Overnight cultures were diluted to $OD_{620}$=0.003 in fresh CDM with indicated concentrations of ET. For endpoint assays, early-log cells ($OD_{620}$=0.2) were collected and washed with chilled 1X PBS to remove excess extracellular thiols. For transport activity assays, cells were grown to early log ($OD_{620}$=0.1) and ET was added to the media. Cells were collected at timepoints of 5, 10, 15, 30, 45, and 60 minutes. 1 mL of each growth was collected for protein quantification using a standard Bradford Assay and the other 4 mL was used for thiol quantification using electrophile capping strategy and isotopically heavy ($d_4$) and natural abundance β-(4-hydroxyphenyl)ethyl iodoacetamide (HPE-IAM) (*24*) essentially as described (*1*). Cells for thiol quantification were resuspended in labeling buffer (5 mM HPE-IAM, 1% DMSO) and subjected to multiple freeze-thaw cycles in liquid nitrogen and a 37°C water bath. The final thaw cycle was for 1 h to allow sufficient alkylation of all thiol containing compounds. Cell lysates were filtered through a 0.22 mm spin filter prior to analysis by isotope-dilution mass spectrometry essentially as described earlier (*1*). Intracellular ergothioneine content estimates were calculated using a value of $5.5x10^8$ cells per unit $OD_{620}$ with an intracellular volume of $6x10^{-16}$ L per cell (*25, 26*). We note that these estimates are approximate and assume identical cell-associated volume and average chain length of two cells for all pneumococcal strains, independent of the growth media, and does not distinguish between cytoplasmic or externally bound ET.

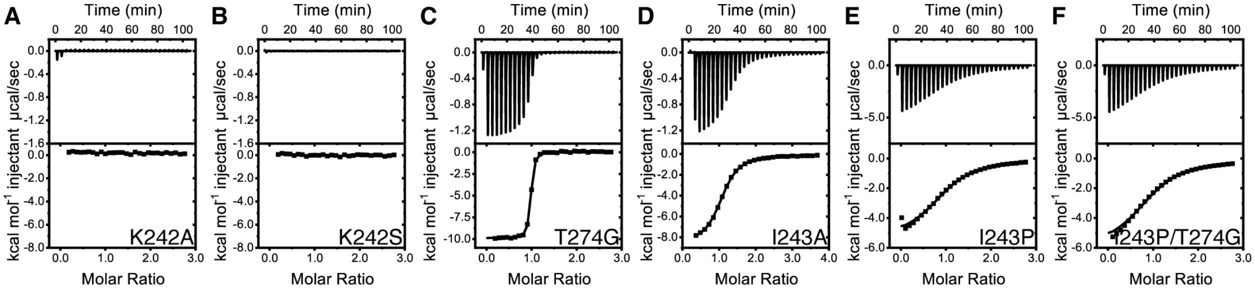


**Fig. S1.**

Representative isothermal titration calorimetry curves of mutant *Sp*EgtUCs with ET. (**A**) K242A (**B**) K242S (**C**) T274G (**D**) I243A (**E**) I243P and (**F**) I243P/T274G SpEgtUC. 20 µM protein was used for titrations shown in panels A-D, with 200 µM protein used for panels E and F Experimental conditions of 50 mM HEPES, pH 7.5, 150 mM NaCl, 2 mM EDTA, 25 °C. Thermodynamic parameters from three replicate experiments are compiled in Fig. 1D and Table 1 (main text).

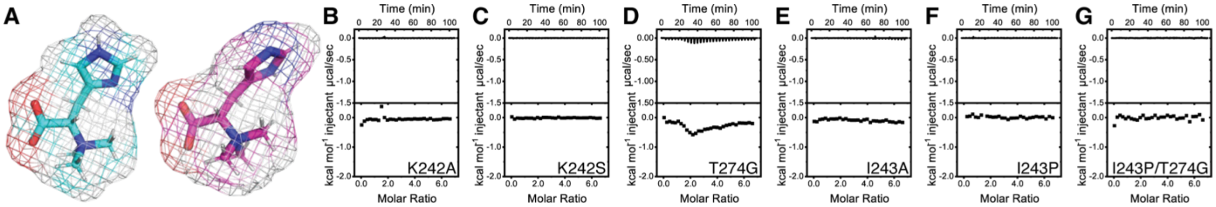


**Fig. S2.**

EgtUC mutants do not bind hercynine (HER). (A) Structures of the two tautomers of hercynine. Representative ITC titrations of 20 µM EgtUC mutants (**B**) K242A (**C**) K242S (**D**) T274G (**E**) I243A (**F**) I243P and (**G**) I243P/T274G titrated with 600 µM HER. Experimental conditions of 50 mM HEPES, pH 7.5, 150 mM NaCl, 2 mM EDTA, 25°C. These data could not be quantitatively analyzed as there is no appreciable heat of binding in any case. NMR- and fluorescence-based assays reveal that HER binds to the wild-type domain with $K_a \approx 10^3$ $M^{-1}$ (*1*) and thus would not be detected under these solution conditions.

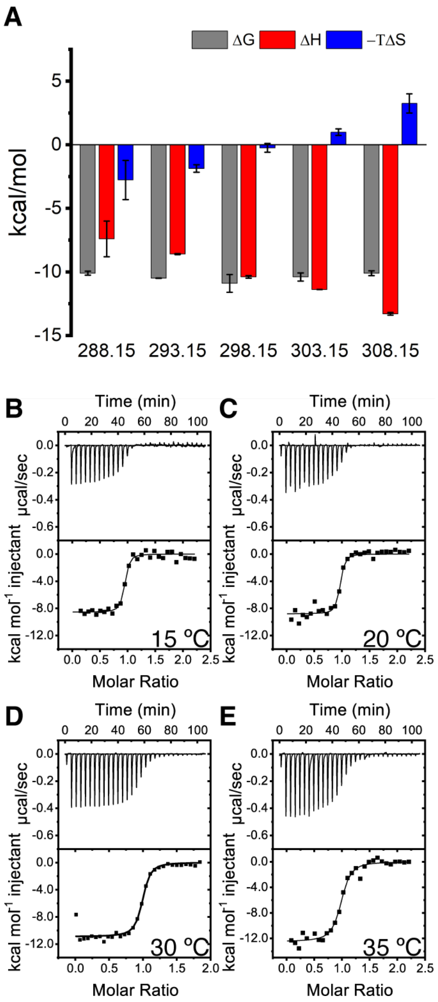


**Fig. S3**

Temperature (*T*)-dependence (*n*=3) of the thermodynamics of ET binding to SpEgtUC. (**A**) Bar graph representation of the *T*-dependence illustrating enthalpic (Δ*H*) and entropic (*T*Δ*S*) contributions of the free energy (Δ*G*) of binding at each temperature. (**B**) Representative ITC interferograms for the binding of ET to *Sp*EgtUC at the indicated temperature. Refers to **Figure 1E**, main text.

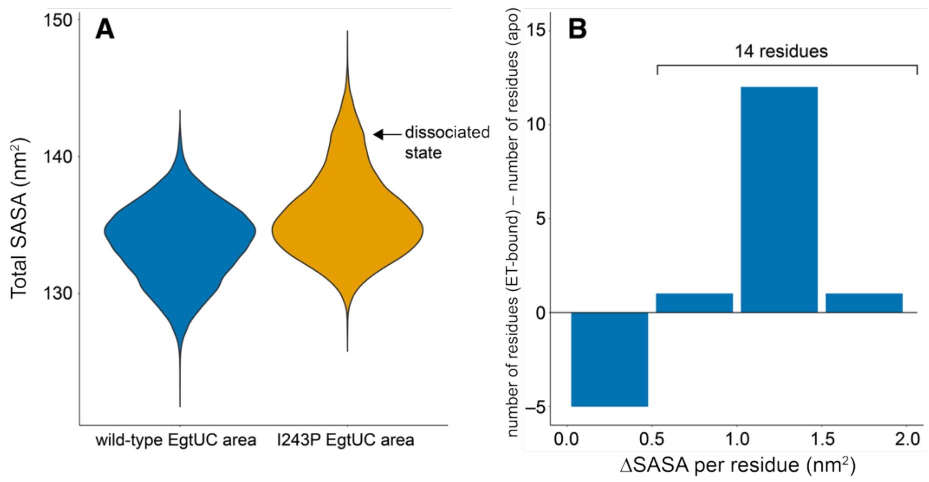


**Fig. S4.**

Solvent accessible surface area of *Sp*EgtUC. (**A**) Solvent accessible surface area of WT and I243P *Sp*EgtUC calculated by molecular dynamics simulations (see Fig. 4, main text). (**B**) Histogram plot of the change in solvent accessible surface area (SASA) per residue between ET-bound and unbound states of WT *Sp*EgtUC as a function of area ($nm^2$).

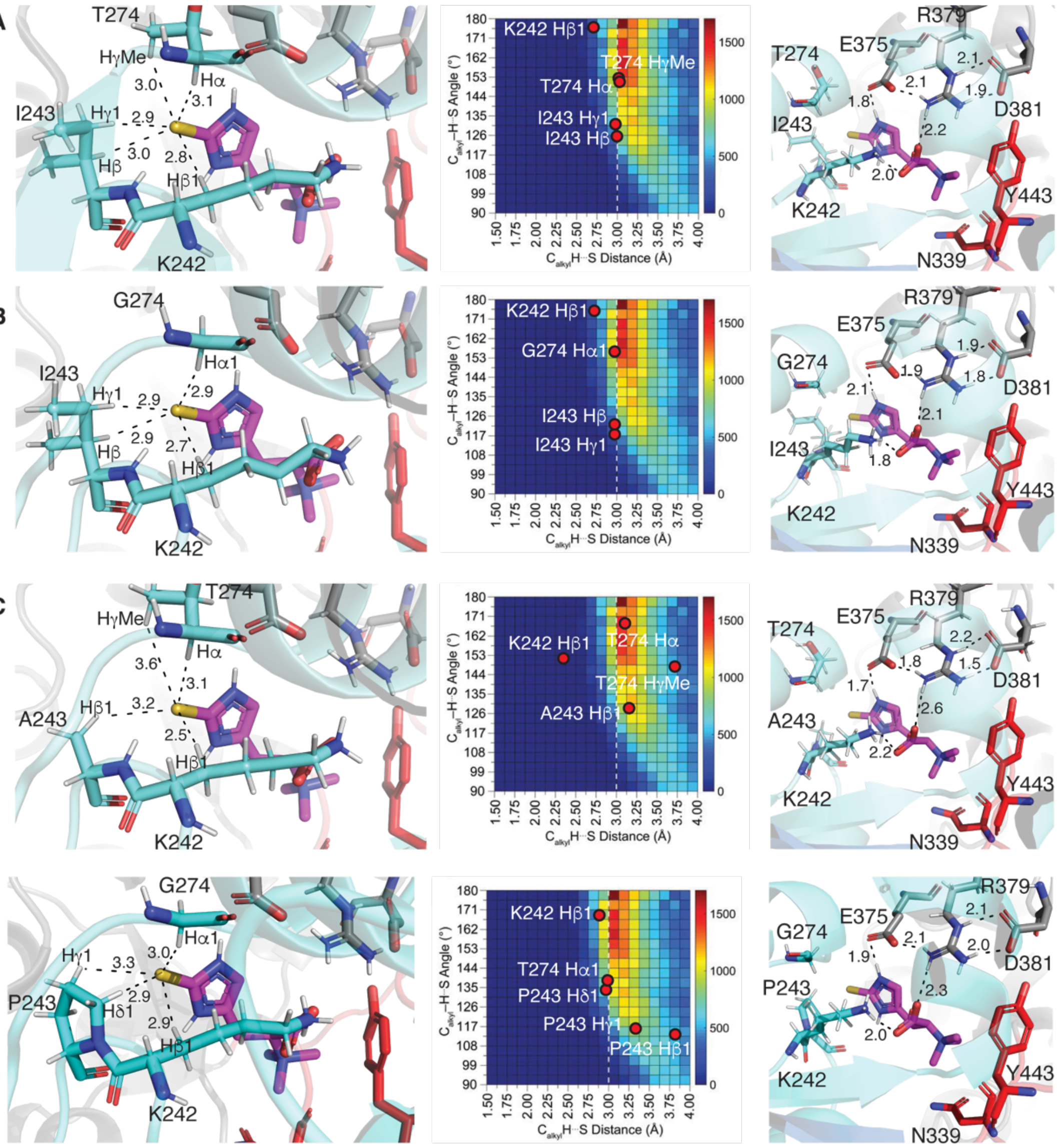


**Fig. S5.**

Crystallographic analysis of various EgtUC domains. *Left to right*, C-H•••S H-bonding interactions with H-S bond lengths noted, heatmap of a histogram of alkyl C-H•••S H-bond lengths and angles from an analysis of the Cambridge Structural Database (CSD) reproduced from reference (*27*), with indicated CH•••S H-bonds for structure indicated, and N-H•••O hydrogen bonding interactions in (**A**) wild-type *Sp*EgtUC, (**B**) T274G *Sp*EgtUC, (**C**) I243A *Sp*EgtUC, and (**D**) I243P/T274G *Sp*EgtUC.

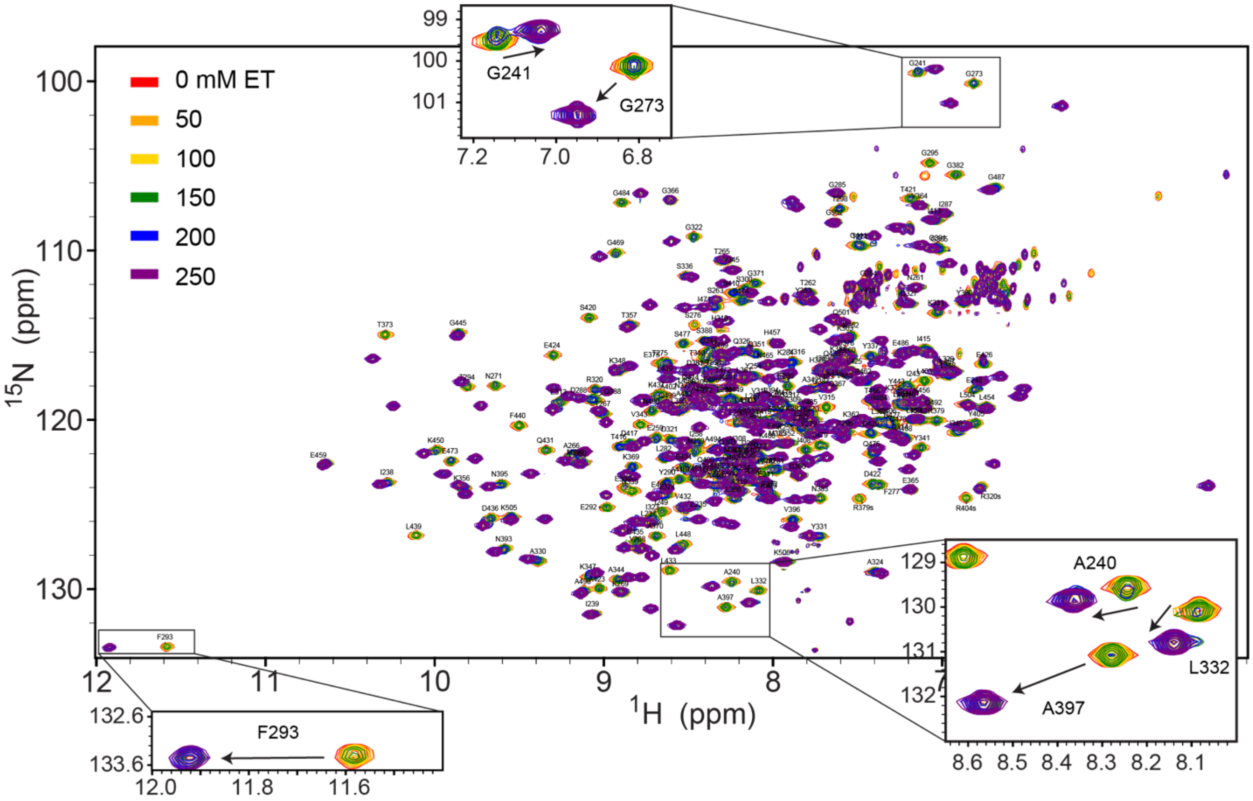


**Fig. S6.**

Full $^{1}$H,$^{15}$N TROSY spectra (800 MHz; 35 $^{o}$C) of 0.2 mM wild-type *Sp*EgtUC without (*red* contours) and with the indicated concentrations of ET added. Intermediate ET concentrations are 50 µM (*orange*), 100 µM (*yellow*), 150 µM (*green*), 200 µM (*blue*) and 250 µM (*purple*). Selected regions are expanded for comparison to the spectra of I243A and I243P shown in **Figure 3**, main text.

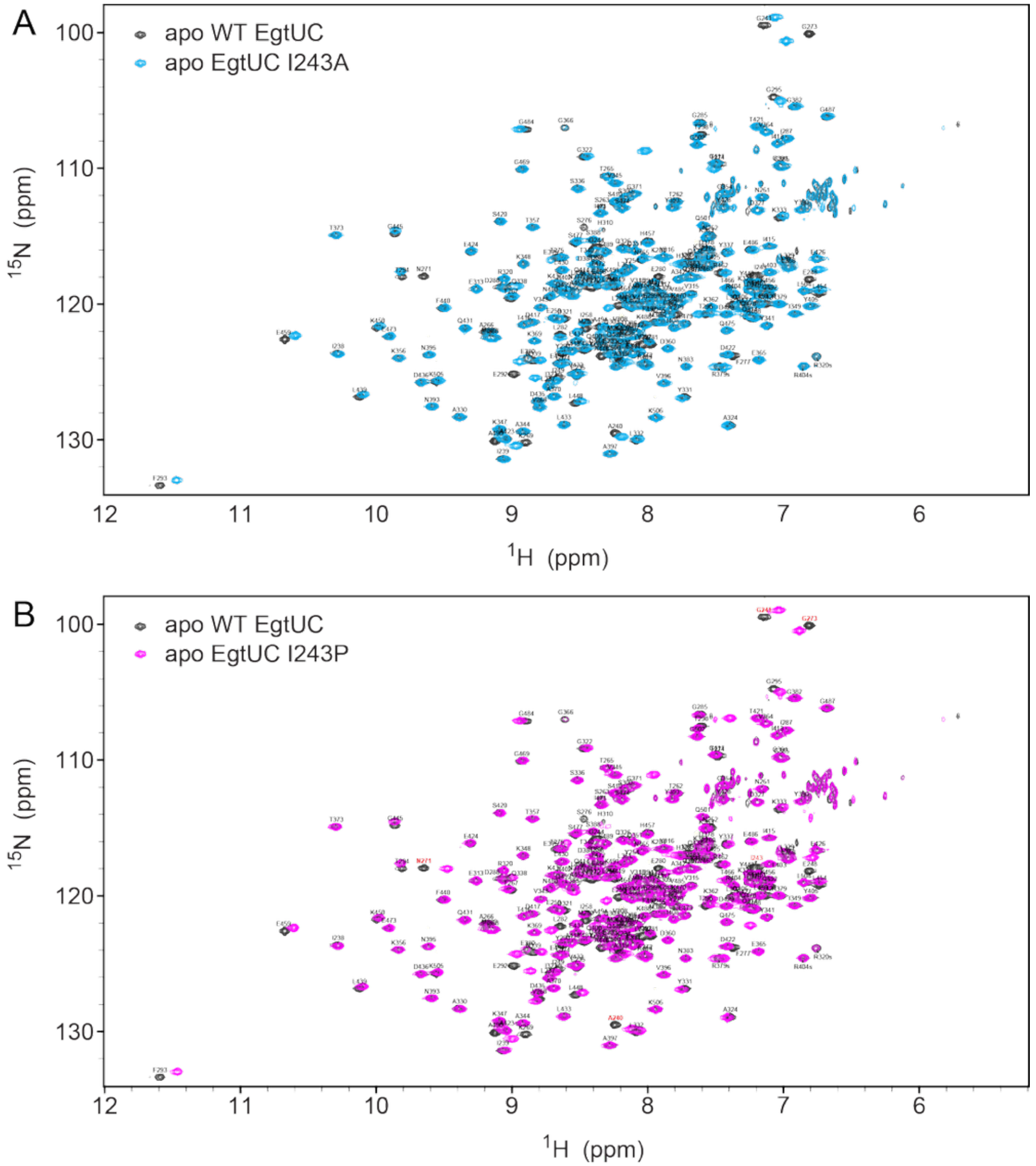


**Fig. S7.**
Full $^{1}$H,$^{15}$N TROSY spectra (600 MHz; 35 $^{o}$C) of apo WT (*black* contours) and (**A**) apo I243A (*blue*) and (**B**) apo WT (*black*) and apo I243P (*magenta*). WT crosspeak assignments (*1*) are provided. These spectra reveal that the ligand-free states of the both I243 substitution mutants are structurally wild-type like, with chemical shift perturbations observed localized to the site of the mutation.

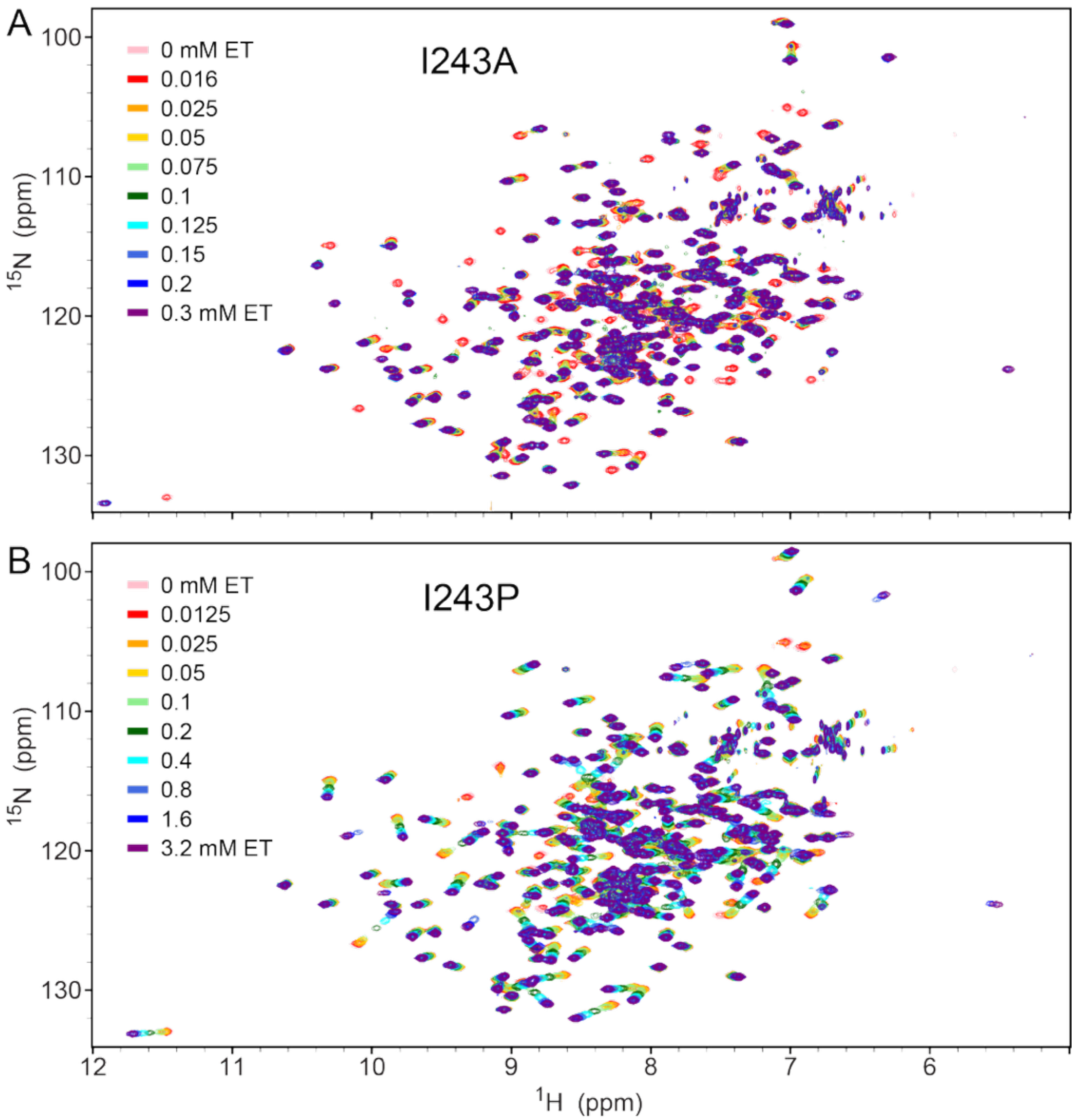


**Fig. S8.**

Full $^{1}H$,$^{15}N$ TROSY spectra (600 MHz; 35 °C) of (**A**) 0.1 mM I243A *Sp*EgtUC without and with the indicated concentrations of ET added and of (**B**) 0.4 mM I243P *Sp*EgtUC without and with the indicated concentrations of the ET added.

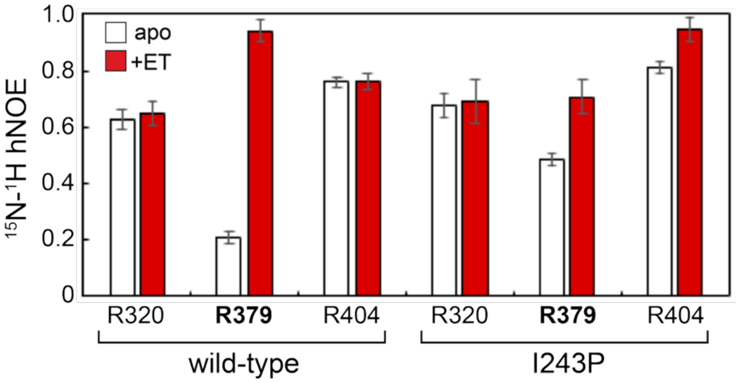


**Fig. S9.**
The value of the $^{15}N$-$^{1}H$-hNOE for the $^{15}N^{e}$-$^{1}H^{e}$ pair for each of the three Arg residues in WT *Sp*EgtUC (*1*) and I243P *Sp*EgtUC in the ligand-free apo (open bars) and ET-bound (closed bars) structures. The error bars incorporate the signal-to-noise in these spectra.

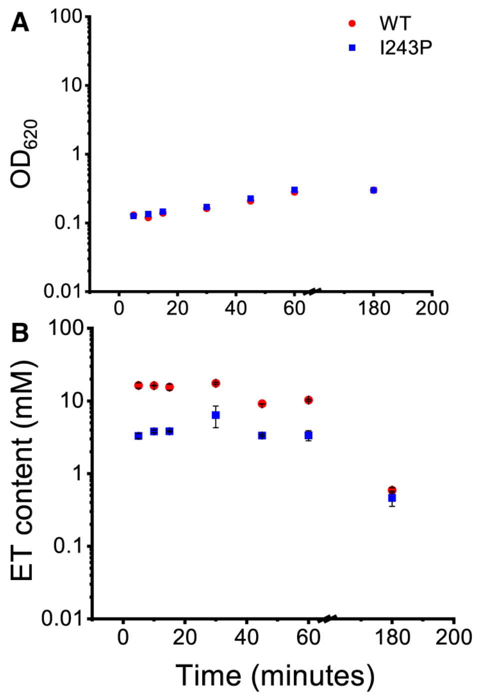


**Fig. S10.**

The rate of ergothioneine import by *Streptococcus pneumoniae* is dependent on the $K_d$ of the *Sp*EgtUC for ET. (**A**) The $OD_{620}$ of cells collected at each timepoint for WT (red circles) and I243P (blue squares). (**B**) ET content in *S. pneumoniae* cells WT (red circles) and I243P (blue squares) quantified by LC-MS at six timepoints, with ET added to a final concentration of 0.05 μM at *t*=0. Data are representative of technical duplicates, with thiol content normalized to cell density. Refers to **Figure 5B**, main text.

**Table S1.**

Thermodynamic parameters obtained for the temperature dependence of ET binding *Sp*EgtUC.

| Temperature (°C) | Method | $K_a$ ($M^{-1}$) | $\Delta G$ (kcal $mol^{-1}$) | $\Delta H$ (kcal $mol^{-1}$) | $-T\Delta S$ (kcal $mol^{-1}$) | *n* |
|---|---|---|---|---|---|---|
| 15 | ITC* | $5.0 \pm 1.4 \times 10^7$ | –10.1 ± 0.1 | –7.4 ± 1.6 | -2.8 ± 1.5 | 1.0 ± 0.2 |
| 20 | ITC | $6.6 \pm 0.2 \times 10^7$ | –10.5 ± 0.1 | –8.6 ± 0.2 | -1.9 ± 0.2 | 0.96 ± 0.04 |
| 25 (*1*) ‡ | ITC | $1.7 \pm 0.1 \times 10^7$ | –10.9 ± 0.1 | –10.4 ± 0.1 | 0.5 ± 0.1 | 0.99 ± 0.01 |
| 30 | ITC | $3.1 \pm 1.3 \times 10^5$ | –10.4 ± 0.2 | –11.4 ± 0.01 | 1.0 ± 0.3 | 0.94 ± 0.01 |
| 35 | ITC | $1.4 \pm 0.4 \times 10^4$ | –10.1 ± 0.1 | –13.3 ± 0.7 | 3.2 ± 0.8 | 0.93 ± 0.1 |

*Solution conditions: 50 mM HEPES, pH 7.5, 150 mM NaCl, 2 mM EDTA, 25.0 ºC. Data are the average and standard deviation of 3 replicates.

‡Reported in ref (*1*). Solution conditions: 50 mM HEPES, pH 7.5, 150 mM NaCl, 2 mM EDTA, 25.0 ºC.

**Table S2.**

Crystallographic statistics for the mutant *Sp*EgtUC structures

| | EgtUC I243A | EgtUC T274G | EgtUC I243P | EgtUC I243P T274G |
|---|---|---|---|---|
| *Data collection* | | | | |
| Wavelength (Å) | 1.00002 | 1.00002 | 1.00002 | 1.00002 |
| Space group | F 2 2 2 | F 2 2 2 | C2 | I41 2 2 |
| *Cell dimensions* | | | | |
| a, b, c (Å) | 118.16 129.20 206.46 | 120.15 127.31 206.65 | 119.75 209.43.72 87.97 | 87.85 87.85 207.64 |
| α, β, γ (°) | 90.00 90.00 90.00 | 90.00 90.00 90.00 | 90.00 132.88 90.00 | 90.00 90.00 90.00 |
| Resolution (Å) | 80.33 – 2.14 (2.49 – 2.14) | 51.94 – 1.95 (2.22 – 1.95) | 64.47 – 1.89 (1.93 – 1.89) | 80.91 – 1.36 (1.39 – 1.36) |
| *Ellipsoidal Resolution* (Å) | | | | |
| a* | 2.92 (a*) | 2.45 (a*) | N/A | N/A |
| b* | 2.61 (b*) | 2.36 (b*) | N/A | N/A |
| c* | 2.14 (c*) | 1.95 (c*) | N/A | N/A |
| $R_{sym}$ | 0.107 (0.290) | 0.108 (0.991) | 0.061 (0.901) | 0.069 (1.866) |
| $R_{meas}$ | 0.119 (0.363) | 0.118 (1.109) | 0.072 (1.069) | 0.072 (1.975) |
| $R_{pim}$ | 0.051 (0.213) | 0.046 (0.489) | 0.039 (0.570) | 0.020 (0.632) |
| Total reflections | 115662 (2950) | 214976 (8197) | 429514 (21739) | 1088844 (36910) |
| No. unique reflections | 23521 (1176) | 33902 (1696) | 124653 (6288) | 85513 (4041) |
| CC1/2 | 0.996 (0.865) | 0.998 (0.652) | 0.999 (0.561) | 1.000 (0.425) |
| I/σ(I) | 8.3 (2.3) | 8.8 (1.8) | 13.1 (1.3) | 23.8 (1.2) |
| Completeness spherical (%) | 53.8 (7.3) | 58.7 (9.2) | 99.3 (99.8) | 99.1 (94.9) |
| Completeness ellipsoidal (%) | 88.8 (52.4) | 88.0 (56.7) | N/A | N/A |
| Multiplicity | 4.9 (2.5) | 6.3 (4.8) | 3.4 (3.5) | 12.7 (9.1) |
| Wilson B-factor | 31.20 | 34.78 | 29.71 | 16.33 |
| *Refinement* | | | | |
| Resolution (Å) | 80.33 – 2.14 (2.23 – 2.14) | 51.94 – 1.95 (2.01 – 1.95) | 64.47 – 1.89 (1.96 – 1.89) | 80.91 – 1.36 (1.41 – 1.36) |
| No. unique reflections | 23513 | 33796 | 124582 | 85298 (8202) |
| $R_{work}$ | 0.2263 | 0.2005 | 0.2137 | 0.1442 |
| $R_{free}$ | 0.2682 | 0.2434 | 0.2542 | 0.1673 |
| *R.m.s.d values* | | | | |
| Bond lengths (Å) | 0.005 | 0.008 | 0.005 | 0.009 |
| Bond angles (°) | 0.860 | 1.089 | 0.768 | 1.054 |
| *No. atoms* | | | | |
| Protein | 4292 | 4300 | 8627 | 2217 |
| Ligand/ions | 30 | 30 | 75 | 53 |
| solvent | 228 | 261 | 1384 | 428 |
| *B-factors* (Å$^2$) | | | | |

| | | | | |
|---|---|---|---|---|
| Protein | 40.51 | 44.07 | 35.36 | 24.38 |
| ligand | 32.93 | 40.20 | 27.60 | 44.24 |
| solvent | 33.77 | 43.60 | 42.78 | 41.88 |
| *Ramachandran plot* | | | | |
| Favored (%) | 98.3 | 98.5 | 98.4 | 98.2 |
| Allowed (%) | 1.3 | 1.1 | 1.2 | 1.4 |
| Outliers (%) | 0.4 | 0.4 | 0.4 | 0.4 |
| Clashscore | 3.58 | 8.89 | 5.92 | 2.01 |
| Rotamer outliers (%) | 0.64 | 0.43 | 0.00 | 0.00 |
| *PDB code* | 9EGH | 9EGI | 9EGJ | 9O5F |

**Table S3.**
*Streptococcus pneumoniae* strains used in this work

| Strain Number | Genotype (Description) | Antibiotic Resistance | Reference or source |
|---|---|---|---|
| IU1781 | D39 *rpsL1* | $Str^R$ | (*1*) |
| IU18400 | D39 *rpsL1 ΔegtUBC::PckanrpsL* | $Kan^R$ | (*1*) |
| IU20937 | D39 *rpsL1 egtUBC*(I243A) | $Str^R$ | This study |
| IU20940 | D39 *rpsL1 egtUBC*(I243P) | $Str^R$ | This study |
| IU20942 | D39 *rpsL1 egtUBC*(T274G) | $Str^R$ | This study |

**Table S4.**
Strain construction primers used in this study

| Primer Name | Primer Sequence (5'-3') | Description |
|---|---|---|
| KL039 | GGAAAACTTGATTATTGCTGGGAAAgcAGGCCCAGAACCAGAAATTTTGG | IU20937: *Sp*EgtUBC I243A FP |
| KL041 | GGAAAACTTGATTATTGCTGGGAAAccAGGCCCAGAACCAGAAATTTTGG | IU20940: *Sp*EgtUBC I243P FP |
| KL042 | CCAAAATTTCTGGTTCTGGGCCTGGTTTCCCAGCAATAATCAAGTTTTCC | IU20940: *Sp*EgtUBC I243P RP |
| KL047 | GTTAAACCGAATTTTGGGGGGACAAGTTTCCTTTATGAAGCTCTGAAAAAAGGTG | IU20942: *Sp*EgtUBC T274G FP |
| KL048 | CACCTTTTTTCAGAGCTTCATAAAGGAAACTTGTCCCCCCAAAATTCGGTTTAAC | IU20942: *Sp*EgtUBC T274G RP |
| 1642kanrpsL_OP1 | GCTGAATATAGTGTCACTTTTGACTTTGTTTTCC | *Sp*EgtUC outside amplification FP |
| KL066 | ACCGATATTGCTCATCAGCTGGCCATTTCAACTTCAACTGTCATTCG | *Sp*EgtUC outside amplification RP |
| KL067 | GCATCGACTTCCAAGTGAGCTATCTGGTGGAGAACAGCAACGG | *Sp*EgtUBC sequencing FP |
| KL068 | AGGACAAAGAGAAAATCGTCAACGCCCTTCAACTACCCTATTCAAACGCC | *Sp*EgtUBC sequencing RP |
| KL075 | GGATTGCCTTTGGGATGACCAGATG | *Sp*EgtUBC intergene sequencing primer |
| KL076 | CCAAAATTTCTGGTTCTGGGCCTGCTTTCCCAGCAATAATCAAGTTTTCCTTCTC | IU20937: *Sp*EgtUBC I243A RP |

**Table S5.**
Plasmid construction primers used in this study

| Primer Name | Primer Sequence (5’-3’) | Description |
|---|---|---|
| KL043 | GGAAAACTTGATTATTGCTGGGGCAATAGGCCCAGAACCA GAAATTTTGG | pSUMO-*Sp*EgtUC K242A FP |
| KL044 | CCAAAATTTCTGGTTCTGGGCCTATTGCCCCAGCAATAATC AAGTTTTCC | pSUMO-*Sp*EgtUC K242A RP |
| KL045 | GGAAAACTTGATTATTGCTGGGAGCATAGGCCCAGAACCA GAAATTTTGG | pSUMO-*Sp*EgtUC K242S FP |
| KL046 | CCAAAATTTCTGGTTCTGGGCCTATGCTCCCAGCAATAATC AAGTTTTCC | pSUMO-*Sp*EgtUC K242S RP |
| KL047 | GTTAAACCGAATTTTGGGGGGACAAGTTTCCTTTATGAAGC TCTGAAAAAAGGTG | pSUMO-*Sp*EgtUC T274G FP |
| KL048 | CACCTTTTTTCAGAGCTTCATAAAGGAAACTTGTCCCCCCA AAATTCGGTTTAAC | pSUMO-*Sp*EgtUC T274G RP |
| KL061 | TGCTGGGAAAgcgGGCCCAGAAC | pSUMO-*Sp*EgtUC I243A FP |
| KL062 | ATAATCAAGTTTTCCTTCTCC | pSUMO-*Sp*EgtUC I243A and I243P RP |
| KL063 | TGCTGGGAAAccgGGCCCAGAAC | pSUMO-*Sp*EgtUC I243P FP |
| KL102 | GTTTTCCTTCTCCATATGGGATCCTCCAATCTGTTC | pSUMO-*Sp*EgtUC backbone amplification RP |
| KL103 | GTCATGAGCCAGAGCAGGTTTATCAAGTG | pSUMO-*Sp*EgtUC backbone amplification FP |
| KL104 | GATCCCATATGGAGAAGGAAAACTTGATTATTGCTGG | pSUMO-*Sp*EgtUC I243P/T274G gene-block amplification FP |
| KL105 | AAACCTGCTCTGGCTCATGACTCACCTTG | pSUMO-*Sp*EgtUC I243P/T274G gene-block amplification RP |

**Movie S1.**

1 μs molecular dynamics simulation of wild-type *Sp*EgtUC bound to ergothioneine using pdb code 7TXK (*1*) as the starting coordinates. Domains are colored as indicated in **Figure 1**, main text. Ergothioneine is in gold in stick, with key residues shown in stick. Refers to **Figure 4**, main text.

**Movie S2.**

1 μs molecular dynamics simulation for I243P *Sp*EgtUC bound to ergothioneine using PDB code 9EGJ as the starting coordinates. Domains are colored as indicated in **Figure 1**, main text. Ergothioneine is in gold stick with key residues shown in stick. From 0-660 ns complex [A], closed state [A], 660-860 ns pre-dissociation state, [B] 860 ns–1 μs, dissociated state [C]. Refers to **Figure 4**, main text.

### References


1. Y. Zhang *et al.*, Discovery and structure of a widespread bacterial ABC transporter specific for ergothioneine. *Nat Commun* **13**, 7586 (2022).
2. T. Lundback, S. van Den Berg, T. Hard, Sequence-specific DNA binding by the glucocorticoid receptor DNA-binding domain is linked to a salt-dependent histidine protonation. *Biochemistry* **39**, 8909-8916 (2000).
3. A. Paleskava, A. L. Konevega, M. V. Rodnina, Thermodynamics of the GTP-GDP-operated conformational switch of selenocysteine-specific translation factor SelB. *J Biol Chem* **287**, 27906-27912 (2012).
4. W. Kabsch, Xds. *Acta Crystallogr D Biol Crystallogr* **66**, 125-132 (2010).
5. I. J. Tickle *et al.* (Global Phasing Ltd., Cambridge, UK, 2018).
6. P. V. Afonine *et al.*, Towards automated crystallographic structure refinement with phenix.refine. *Acta Crystallogr D Biol Crystallogr* **68**, 352-367 (2012).
7. P. Emsley, K. Cowtan, Coot: model-building tools for molecular graphics. *Acta Crystallogr D Biol Crystallogr* **60**, 2126-2132 (2004).
8. V. B. Chen *et al.*, MolProbity: all-atom structure validation for macromolecular crystallography. *Acta Crystallogr D Biol Crystallogr* **66**, 12-21 (2010).
9. M. J. Robertson, J. Tirado-Rives, W. L. Jorgensen, Improved Peptide and Protein Torsional Energetics with the OPLSAA Force Field. *J Chem Theory Comput* **11**, 3499-3509 (2015).
10. H. J. C. Berendsen, J. R. Grigera, T. P. Straatsma, The missing term in effective pair potentials. *J. Phys. Chem.* **91**, 6269–6271 (1987).
11. L. S. Dodda, I. Cabeza de Vaca, J. Tirado-Rives, W. L. Jorgensen, LigParGen web server: An automatic OPLS-AA parameter generator for organic ligands. *Nucleic Acids Res* **45**, W331-W336 (2017).
12. M. J. Abraham *et al.*, GROMACS: High performance molecular simulations through multi-level parallelism from laptops to supercomputers. *SoftwareX* **1-2**, 19-25 (2015).
13. S. Pronk *et al.*, GROMACS 4.5: A high-throughput and highly parallel open source molecular simulation toolkit. *Bioinformatics* **29**, 845-854 (2013).

14. B. Hess, C. Kutzner, D. van der Spoel, E. Lindahl, GROMACS 4: Algorithms for highly Efficient, load-balanced, and scalable molecular simulation. *J Chem Theory Comput* **4**, 435-447 (2008).
15. B. Hess, H. Bekker, H. J. C. Berendsen, J. G. E. M. Fraaije, LINCS: A Linear Constraint Solver for molecular simulations. *J. Comp. Chem.* **18**, 1463-1472 (1998).
16. S. Miyamoto, P. A. Kollman, Settle: An analytical version of the SHAKE and RATTLE algorithm for rigid water models. *J. Comp. Chem.* **13**, 952-962 (1992).
17. U. Essmann *et al.*, A smooth particle mesh Ewald method. *J. Chem. Phys.* **103**, 8577-8592 (1995).
18. G. Bussi, D. Donadio, M. Parrinello, Canonical sampling through velocity rescaling. *J. Chem. Phys.* **126**, 014101 (2007).
19. M. Parrinello, A. Rahman, Polymorphic transitions in single crystals: A new molecular dynamics method *J. Appl. Phys.* **52**, 7182-7190 (1981).
20. E. C. Meng *et al.*, UCSF ChimeraX: Tools for structure building and analysis. *Protein Sci* **32**, e4792 (2023).
21. H. Wickham. (Springer-Verlag, New York, 2016).
22. F. Eisenhaber, P. Lijnzaad, P. Argos, C. Sander, M. Scharf, The double cubic lattice method: Efficient approaches to numerical integration of surface area and volume and to dot surface contouring of molecular assemblies. *J. Comp. Chem.* **16**, 273-284 (1995).
23. I. van de Rijn, R. E. Kessler, Growth characteristics of group A streptococci in a new chemically defined medium. *Infect Immun* **27**, 444-448 (1980).
24. H. A. Hamid *et al.*, Polysulfide stabilization by tyrosine and hydroxyphenyl-containing derivatives that is important for a reactive sulfur metabolomics analysis. *Redox Biol* **21**, 101096 (2019).
25. F. E. Jacobsen, K. M. Kazmierczak, J. P. Lisher, M. E. Winkler, D. P. Giedroc, Interplay between manganese and zinc homeostasis in the human pathogen *Streptococcus pneumoniae*. *Metallomics* **3**, 38-41 (2011).
26. S. Ramos-Montanez, K. M. Kazmierczak, K. L. Hentchel, M. E. Winkler, Instability of ackA (acetate kinase) mutations and their effects on acetyl phosphate and ATP amounts in Streptococcus pneumoniae D39. *J, Bacteriol* **192**, 6390-6400 (2010).
27. H. A. Fargher, T. J. Sherbow, M. M. Haley, D. W. Johnson, M. D. Pluth, C-H•••S hydrogen bonding interactions. *Chem Soc Rev* **51**, 1454-1469 (2022).
